# PWO1 and TRB proteins coordinate chromatin regulation to prevent premature differentiation and ectopic lignin deposition in Arabidopsis

**DOI:** 10.64898/2026.07.24.740627

**Authors:** Ahamed Khan, Alžbeta Kusová, Jan Skalák, Biswajit Ghosh, Tingting Yang, Antonia Lilly Vivien Kelling, Laura Hagemann, Kishore C S Panigrahi, Jan Hejátko, Petra Procházková Schrumpfová, Yue Zhou, Sara Farrona, Iva Mozgová, Daniel Schubert

## Abstract

The Arabidopsis PWWP-DOMAIN INTERACTOR OF POLYCOMBS1 (PWO1) and Telomere Repeat-Binding Proteins 1-3 (TRB1-3, TRBs) associate with distinct and shared protein complexes involved in epigenetic regulation, yet their cooperative roles in chromatin control and plant development remain largely unexplored. Here, we show that the interaction between PWO1 and TRBs is evolutionarily conserved. Both PWO1 and TRBs associate with plant telomeres, interact at these regions, and are co-enriched at subsets of interspersed *telo*-box motifs across regulatory regions genome-wide. TRBs facilitate PWO1 binding at shared genomic regions, including *telo*-box motifs. PWO1 and TRBs share a substantial number of genomic targets and preferentially bind chromatin regions associated with transcriptionally active states, whereas TRBs alone associate with repressive marks at thousands of loci. Genetic analyses show that the *pwo1 trb1 trb3* triple mutant displays severe developmental defects, including main stem arrest and early maturation associated with aberrant lignin deposition in interfascicular tissues. In the triple mutant, key enzymes in the lignin biosynthesis pathway are upregulated, indicating that PWO1, TRB1, and TRB3 cooperatively regulate secondary cell wall formation. Together, our findings provide new insights into how PWO1 and TRBs cooperate to regulate chromatin states and orchestrate plant development, highlighting their central role in controlling gene expression programs.

**Significance statement:** This study shows that PWO1 and TRB proteins co-occupy telomeres, including interspersed *telo*-box motifs, to regulate chromatin organization and plant development, particularly ectopic lignin deposition. Our findings reveal how these nuclear protein factors coordinate epigenetic states in *Arabidopsis thaliana*, providing a framework for understanding the control of developmental programs.

Graphical abstract:
Evolutionarily conserved PWO-TRB interactions and their shared roles in chromatin regulation and plant development. *Created with BioRender.com*.

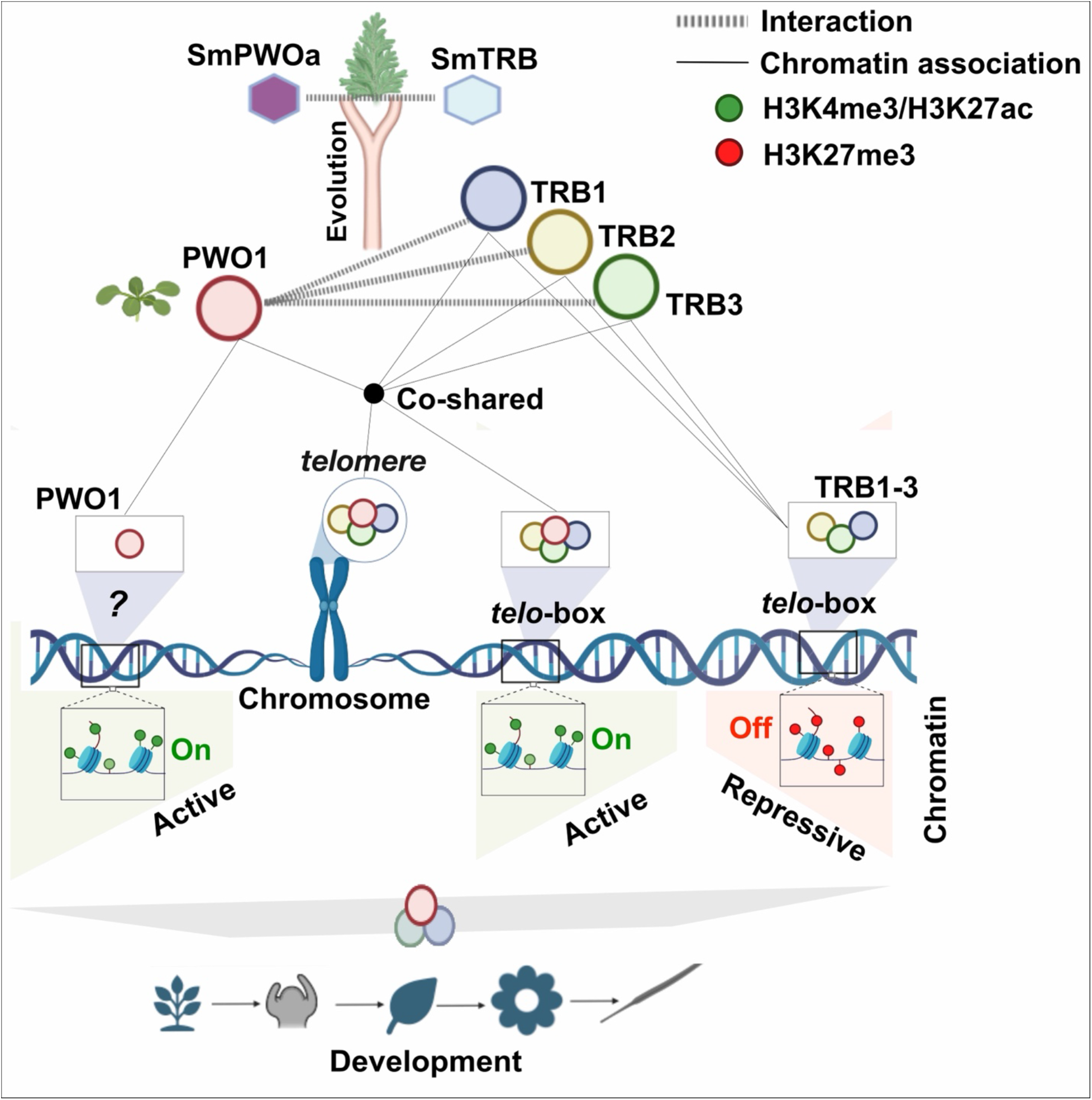

## Introduction

Epigenetic regulators control development in eukaryotes by modifying DNA and histone tails, thereby influencing chromatin architecture and gene accessibility (Hemenway and Gehring 2023). However, most of these proteins (writers and erasers) lack intrinsic DNA-binding domains (Bieluszewski et al. 2021; Wu et al. 2023), and therefore rely on accessory proteins, including readers (Hohenstatt et al. 2018; Zhou et al. 2018; Yuan et al. 2021; Wang et al. 2023), to be recruited to specific chromatin regions for modification of histones or DNA (Scheid et al. 2021). These modifications enable chromatin to locally adopt either active or repressive transcriptional states hence influencing plant development and response to stresses. In this context, plant-specific accessory proteins such as PWWP-DOMAIN INTERACTOR OF POLYCOMBS (PWOs/PWWPs) and Telomere Repeat Binding Proteins (TRBs) are emerging as transcriptional regulators, but their coordinated roles in chromatin dynamics and plant development remain largely unexplored.

Recent studies reveal that *telo-*box motif functions as a gene regulatory hub, recruiting crucial activating or repressing complexes to precisely regulate specific sets of target genes (Zhou et al., 2018; Zhou et al., 2016; Wang et al., 2023; Zheng et al., 2023). In *Arabidopsis thaliana* (Arabidopsis), TRB1-5 were shown to bind to both *terminal telomeric repeats* (telomeres) and *telo-*boxes; a short telomere-related *cis*-regulatory DNA motif (-TTTAGGG-) (Procházková Schrumpfová et al. 2014; Schrumpfová et al. 2016). Specifically, TRB1-3 (TRBs) play a key role in telomere protection and are crucial for Arabidopsis development. Neither the *trb* single nor double mutants show any visible phenotypic defect compared to wild-type controls. In contrast, triple mutants lacking TRB1, TRB2, and TRB3 activity (*trb1 trb2 trb3*) display severe dwarf phenotypes, including cotyledon curling, slow root and leaf growth, and telomere shortening (Procházková Schrumpfová et al. 2014; Zhou et al. 2018; Wang et al. 2023, 2025). TRBs contain a single MYB-like domain, followed by a histone-like H1/5 domain and a C-terminal coiled-coil (CC) domain. The MYB-like domain binds plant telomeric repeats (TTTAGGG)n, while the histone-like (H1/5) domain primarily facilitates intra- and intermolecular protein interactions, including homodimerization and heterodimerization among the TRBs (Amiard et al., 2024; Procházková Schrumpfová et al., 2014; Wang et al., 2025; Kusová et al., 2023). The CC domain of TRBs interacts with the Polycomb Repressive Complex 2 (PRC2) catalytic subunits CURLY LEAF (CLF) and SWNINGER (SWN) (Zhou et al. 2018). TRB1 protein versions lacking either the MYB or H1/5 domain show reduced chromatin binding compared with wild-type TRB1, with the MYB domain providing the strongest sequence-dependent enrichment. Consistent with this, a MYB-deleted TRB1 failed to complement the *trb1 trb2 trb3* mutant, whereas TRB1-H1/5 restored development to a substantial extent (Wang et al. 2025).

TRBs function in cooperation with other proteins or complexes to regulate plant development and chromatin function. For example, TRBs are genetically associated with LIKE HETEROCHROMATIN PROTEIN 1 (LHP1), a component of PRCs, as combinatorial mutants of *trb1-1, trb3-1* and *lhp1-3* exhibit enhanced phenotypic alterations including earlier flowering and stronger leaf size reduction compared to single mutants (Turck et al. 2007; Derkacheva et al. 2013; Zhou et al. 2016). Interestingly, TRBs perform a dual function in establishing a repressive chromatin state in *Arabidopsis*. First, TRBs recruit the PRC2 catalytic subunits to *telo-*box and related motifs, facilitating the deposition of H3K27me3 and maintaining gene repression at specific target genes (Zhou et al. 2016). Secondly, TRBs interact with JUMONJI14 (JMJ14), which promotes the demethylation of the active chromatin mark H3K4me3 at targeted loci (Wang et al. 2023). More recently, TRBs have also been shown to assemble into higher-order regulatory complexes, including the TRB1/2/3–HELIX–TURN–HELIX PROTEIN COMPLEX (TRHT) and the TRB1/2/3–HISTONE-DEMETHYLASE COMPLEX (TRHD), both of which recognize a common set of genomic targets and are required for JMJ14-mediated H3K4me3 demethylation (Wang et al. 2025). Furthermore, TRBs have also been identified as core components of the PEAT complex, which includes PWOs, Enhancer of Polycomb-Related proteins (EPCRs), and AT-rich interaction domain-containing proteins (ARIDs). Within the PEAT complex, TRBs physically interact with PWO1-2, and together with other PEAT components, they play a crucial role in maintaining heterochromatin silencing through their interaction with histone deacetylases (Tan et al. 2018). Additionally, PEAT was also shown to interact with histone acetyltransferases of the MYST family (HAMs) and UBIQUITIN PROTEASE 5 (UBP5) to induce gene expression by histone 4 (H4) acetylation and H2A deubiquitination (Zheng et al., 2023; Godwin et al., 2024). TRB4 and TRB5 have been also shown to interact with PRC2 components, including the catalytic subunits CLF and SWN as well as the Su(z)12 homologues EMBRYONIC FLOWER 2 (EMF2) and VERNALIZATION 2 (VRN2) (Kusová et al., 2023; Amiard et al., 2024). More recently, TRBs paralogs were shown to be partially redundant yet functionally diversified: TRB1 associates with PEAT and the NuA4 acetylation complex, whereas TRB2 and TRB3 associate with PRC2- regulated targets and bind *telo*-box motifs (Mendler et al. 2026). Altogether, TRB proteins are emerging as key recruiters of both active and repressive chromatin modulators, playing a crucial role in regulating cellular and molecular processes essential for plant development.

In Arabidopsis, three orthologs of the PWWP DOMAIN- DOMAIN INTERACTOR OF POLYCOMBS1-3 (PWO1-3/PWWP1-3) have been identified, and they play a critical role in plant development. PWO single and double mutants display only mild morphological phenotypes. In contrast, the *pwo1 pwo2 pwo3* triple mutant exhibits a seedling-lethal phenotype characterized by meristem arrest, demonstrating the essential role of their combined activity in early development and their functional redundancy (Hohenstatt et al. 2018). Structurally, PWO proteins share a conserved PWWP domain, which is a hallmark of chromatin-associated proteins. Aside from this domain, PWOs lack other structured motifs, except for a recently characterized short α-helix located at the C-terminus, termed the C-motif (Khan et al. 2026). PWO1 has been shown to interact with the catalytic subunits of the PRC2, including CLF and SWN (Hohenstatt et al. 2018), and this interaction appears to be evolutionarily conserved across plant species (Khan et al. 2026). Moreover, PWO1 participates in higher-order chromatin organization by binding to the boundaries of H3K27me3-enriched compartment domains (CDs), where it contributes to maintaining their repressive state and regulate their spatial position in the nucleus (Yang et al. 2024). Thus, the functional implications of the PWO1-TRB association are particularly intriguing, as these proteins interact with distinct chromatin regulatory complexes in a context-dependent manner. Exploring their cooperative roles in chromatin regulation and plant development is essential to understand how the PWO1-TRB module fine-tunes gene expression.

In this study, we investigate PWO1-TRBs-mediated chromatin regulation and its role in plant development. We show that the interaction between PWO1 and TRB proteins is evolutionarily conserved and that PWO1, similarly to TRBs, localizes to telomeric regions, including interspersed *telo*-box motifs. Our analyses further indicate that PWO1 and TRBs share a substantial set of genomic targets, suggesting coordinated binding at both genome-wide and *telo*-box-associated sites. Chromatin profiling suggests that these shared targets are predominantly associated with active chromatin states. Importantly, Chromatin Immunoprecipitation analysis of PWO1-GFP in a *trb1 trb2 trb3* mutant background revealed that PWO1 chromatin binding depends on TRB proteins, indicating that TRBs are required for PWO1 association with chromatin and targeting to *telo*-box motifs. Functional genetic analyses reveal that while single and double PWO1 and TRB1- 3 mutants display only mild phenotypes, higher-order disruption of *pwo1 trb1 trb3* leads to pronounced developmental abnormalities, including altered inflorescence stem development and reduced plant size. These phenotypes are associated with mis-regulation of genes involved in secondary cell wall biosynthesis, including pathways linked to lignin deposition. Collectively, our findings suggest that PWO1, in cooperation with TRB proteins, contributes to proper chromatin organization and is important for normal plant development.

## Results

### PWO and TRB proteins interact in an evolutionarily conserved manner and co-localize at telomeric regions in plant nuclei

To investigate whether the interaction between PWO1 and TRBs is evolutionarily conserved, yeast two-hybrid (Y2H) assays were performed. Consistent with previous reports on the PEAT complex (Zheng et al., 2023; Tan et al., 2018), PWO1 physically interacted with both TRB1 and TRB2 (Figure S1A). PWO and TRB proteins from *Selaginella moellendorffii* (Sm), a lycophyte in which PWO proteins first appeared during plant evolution (Khan et al. 2026), were also tested. Hereafter, *S. moellendorffii* PWO and TRB proteins are referred to as SmPWOs and SmTRBs, respectively. A total of six putative TRB proteins were identified in *S. moellendorffii* (Kusová et al. 2025), among which SmTRB (XP_002972715) was characterized. SmPWOa interacted with SmTRB, and cross-species assays showed that Arabidopsis PWO1 interacted with SmTRB, while SmPWOa interacted with Arabidopsis TRB1 and TRB2 (Figure S1B). AlphaFold3 (AF3; Abramson et al., 2024) predictions revealed full-length structural conservation covering MYB, H1/5, and CC domains, among Arabidopsis TRB1-5 and SmTRB (Figure S1C-H), consistent with previous reports (Kusová et al., 2023; Amiard et al., 2024).

PWO1, SmPWOa, and TRB1-3 formed nuclear speckles when expressed individually in *Nicotiana benthamiana* leaves (Kusová et al. 2023; Khan et al. 2026). To assess whether these proteins occupy the same subnuclear space, co-localization analyses were performed. SmTRB-GFP (Figure S2A) and SmTRB-mCherry (Figure S2B) formed speckles in the nucleoplasm and accumulated in the nucleolus, similar to Arabidopsis TRBs (Kusová et al., 2023). Co-expression of SmTRB-GFP with Arabidopsis TRB1-RFP (Figure S2C), TRB2-RFP (Figure S2D), or TRB3-RFP (Figure S2E) resulted in co-localization of GFP- and RFP-positive speckles in the nucleoplasm. Interestingly, co-expression of PWO1-GFP with TRB1-RFP (Figure 1A), TRB2-RFP (Figure 1B), or TRB3- RFP (Figure 1C) also showed strong co-localization at nuclear speckles. SmPWOa-GFP and SmTRB-mCherry also co-localized (Figure 1D). Cross-species co-expression of SmPWOa-GFP with Arabidopsis TRB1-RFP (Figure 1E), TRB2-RFP (Figure 1F), and TRB3-RFP (Figure 1G) showed co-localization within derived speckles. These results indicate that both PWO and TRBs are likely to share the subnuclear space, and this phenomenon is conserved in plant evolution.

**Figure 1.**
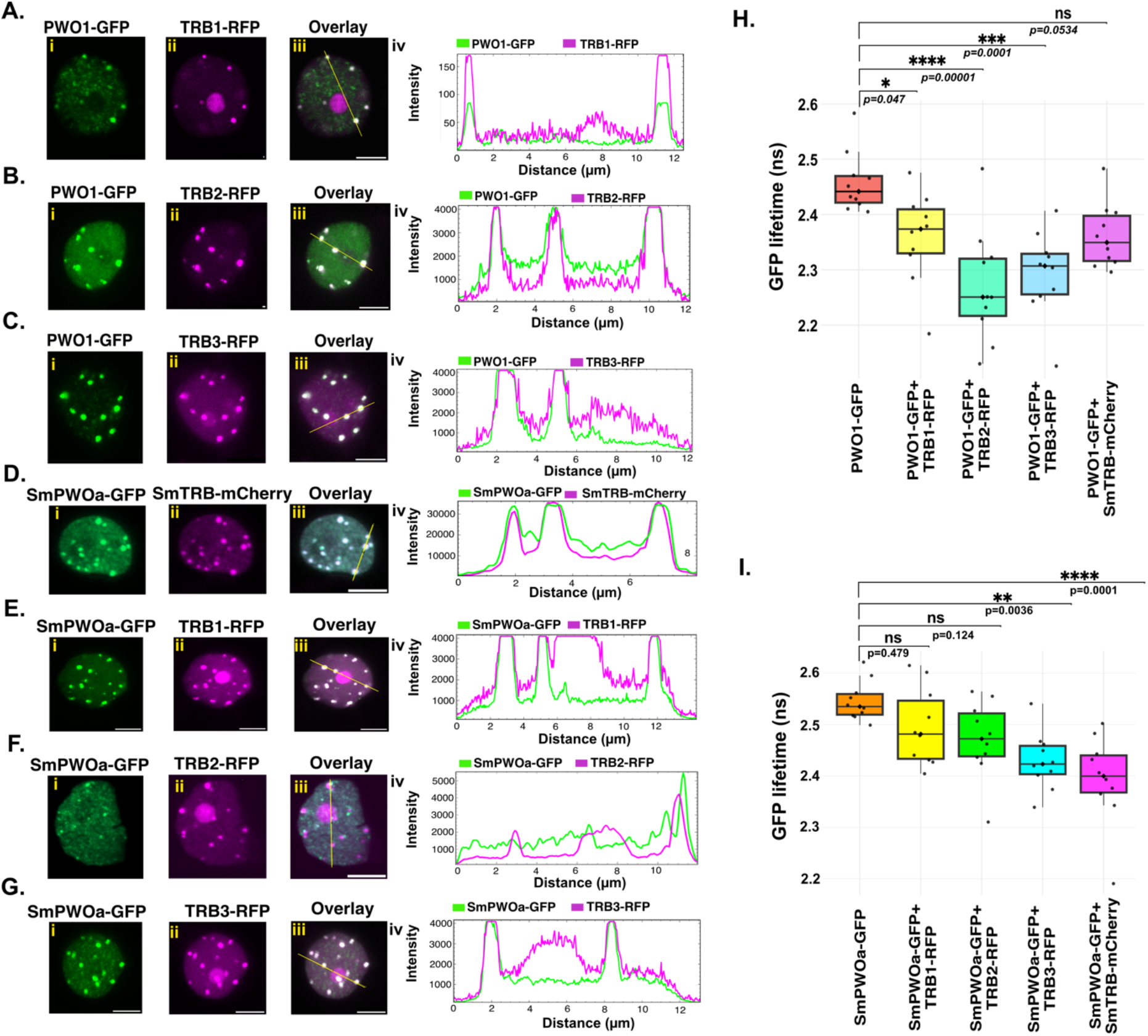
PWO1 and TRBs interact and co-localize at discrete nuclear speckles in *Nicotiana benthamiana* leaf epidermal tissues. Representative confocal microscopy images show nuclei of *N. benthamiana* leaf cells co-infiltrated with *i35S::PWO1-GFP* along with: **A_i–iii**:35S::TRB1-RFP, **B_i–iii**: *35S::TRB2-RFP*, and **C_i–iii**: *35S::TRB3-RFP*. The co-infiltration of *i35S::SmPWOa-GFP* along with **D_i–iii**: *i35S::SmTRB-mCherry*, **E_i–iii**: *35S::TRB1-RFP*, **F_i–iii**: *35S::TRB2-RFP*, and **G_i–iii**: *35S::TRB3-RFP*. For each co-infiltration setup, profiles of GFP and RFP/mCherry fluorescence intensities along the yellow line are shown in **A-G_iv**. Scale bar = 5 µm. FLIM-FRET analyses of GFP lifetime in *N. tabacum* nuclei co-expressing **H.** *i35S::PWO1-GFP* and **I.** *i35S::GFP-SmPWOa* with *35S::TRB1-RFP*, *35S::TRB2-RFP*, *35S::TRB3-RFP*, and *i35S::SmTRB-mCherry* in nuclear condensate (Speckles). Representative images are shown in Figure S2 F-G. Boxes represent interquartile ranges, horizontal lines represent the medians and whiskers represent SE; individual data points are shown. Statistical significance was determined using one-way ANOVA: ****P ≤ 0.0001, ***P ≤ 0.001, **P ≤ 0.01, *P ≤ 0.05. “ns” indicates not significant (P > 0.05).

To test whether PWOs and TRBs interact at nuclear speckles, Förster Resonance Energy Transfer (FRET) with Fluorescence Lifetime Imaging Microscopy (FLIM) was performed in *Nicotiana tabacum* cells. Co-expression of PWO1-GFP with TRB1-RFP, TRB2-RFP or TRB3-RFP significantly reduced the fluorescence lifetime of PWO1-GFP (Figure 1H; Figure S2F), confirming direct interactions within speckles, whereas SmTRB-mCherry showed no significant interaction. Similarly, SmPWOa-GFP showed reduced lifetime when co-expressed with TRB1-3 or SmTRB, with strongest interaction observed for TRB3 and SmTRB within speckles (Figure 1I; Figure S2G). Overall, these findings demonstrated that the interaction between PWO and TRBs is evolutionarily conserved, with both proteins co-localizing and interacting within nuclear speckles.

TRB1-derived speckles have previously been shown to associate with telomeric regions in *N. benthamiana*, based on FISH (Fluorescence *in situ* hybridization) experiments demonstrating telomere co-localization in ∼90% of observed speckles (Schrumpfová et al. 2004). This was further supported by CRISPR-based live imaging of plant telomeres using the dCas9:2xMS2:3xGFP system co-expressed with TRB1-RFP, which confirmed the presence of TRB1 speckles at telomeric sites (Khosravi et al. 2020). Using the same system, dCas9:2xMS2:3xGFP was co-expressed with empty-RFP (Figure 2A), TRB1-RFP (Figure 2B) (Khosravi et al. 2020), TRB2-RFP (Figure 2C) and TRB3-RFP (Figure 2D) showing co-localization between telomeres and TRB-derived speckles (Figure 2B-D). Given that *S. moellendorffii* possesses the canonical TTTAGGG-type telomeres (Shakirov and Shippen 2012), SmTRBs were hypothesized to localize to telomeres in *N. tabacum*, which was confirmed by co-expression of SmTRBs with the dCas9:2xMS2:3xGFP (Figure S2H). Interestingly, PWO1-mCherry also co-localizes with dCas9:2xMS2:3xGFP-labeled telomeres when co-expressed (Figure 2E), indicating that PWO1, like the TRBs, associates with telomeric regions in *planta*. This observation is further supported by FLIM-FRET analyses, where reduced GFP fluorescence lifetimes are detected for dCas9:2xMS2:3xGFP, co-expressed with TRB1-3-(RFP) and PWO1-mCherry compared to the dCas9:2xMS2:3xGFP control within speckles (Figure 2F-G), indicating that TRB1-3 and PWO1 are in close proximity to telomeric DNA in plant nuclei.

**Figure 2:**
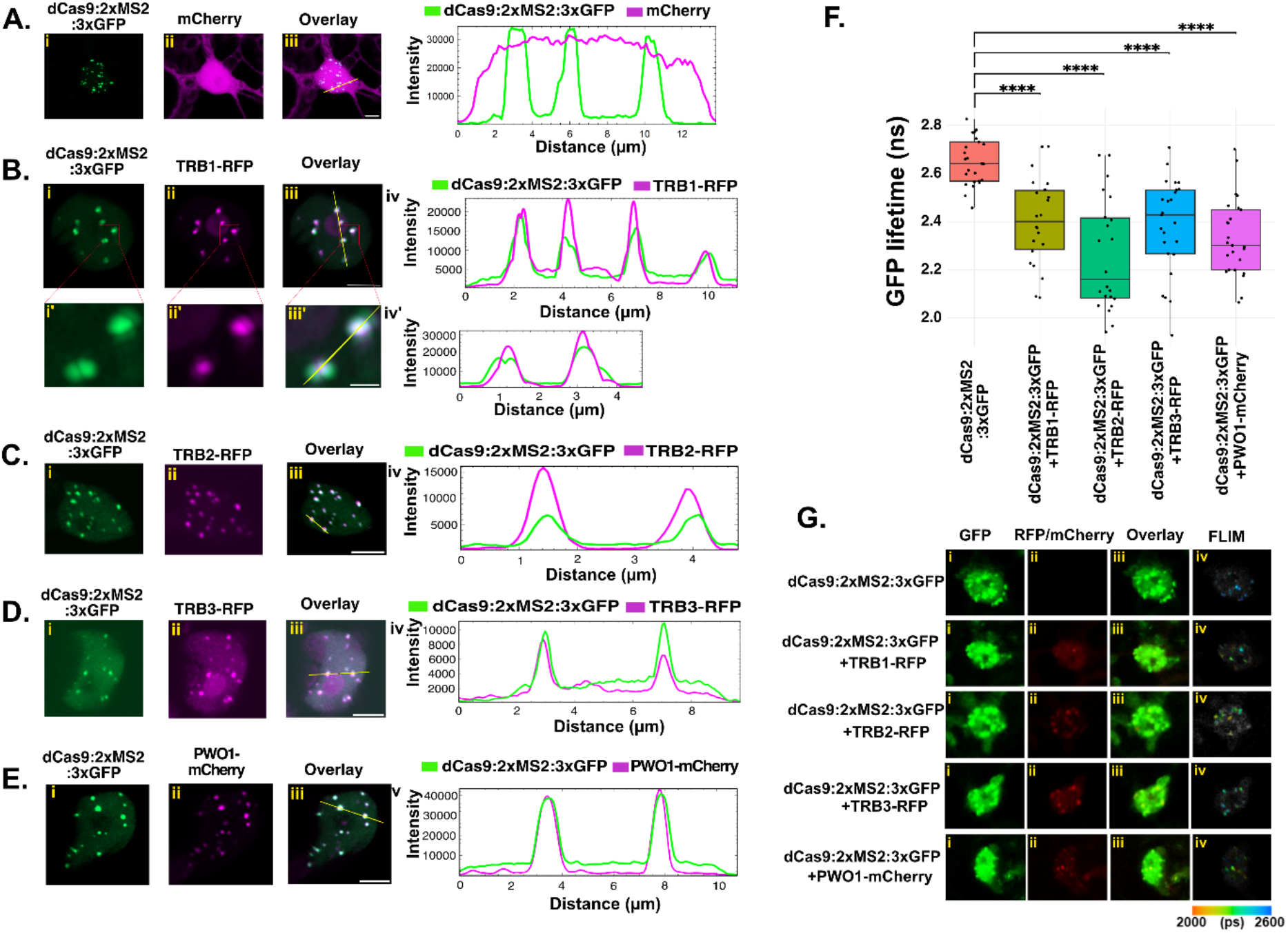
PWO1 and TRBs localization at telomeric regions. Representative confocal microscopy images show the nuclei of *N. benthamiana* leaf cells infiltrated with *dCas9:2XMS2:3XGFP* with **A_i–iii:** RFP, **B_i–iii:** *35S::TRB1-RFP*, including a zoomed-in view **B_i’-iii’** of the telomere pairs closely placed in nucleoplasmic space. Similarly, co-infiltration of *dCas9:2xMS2:3XGFP* along with **C_i–iii:** *35S::TRB2-RFP*, **D_i–iii:** *35S::TRB3-RFP* and **E_i–iii:** *i35S::PWO1-mCherry*. For each co-infiltration setup, profiles of GFP and RFP/mCherry fluorescence intensities along the yellow line are shown in **A–E_iv** and **A_iv’**. Scale bar = 5 µm. For panel **A_i’-iii’**, the scale bar is 2 µm. FLIM-FRET analyses of GFP lifetime in *N. tabacum* nuclei expressing **F.** *dCas9:2xMS2:3XGFP* alone and co-expressing with *35S::TRB1-RFP, 35S::TRB2-RFP, 35S::TRB3-RFP* and *i35S::PWO1-mCherry* in nuclear condensate (speckles). Boxes represent interquartile ranges; horizontal lines represent the medians and whiskers represent SE; individual data points are shown. Statistical significance was determined using one-way ANOVA: ****P ≤ 0.0001. **G.** Representative confocal images showing subnuclear localization of *dCas9:2XMS2:3XGFP with 35S::TRB1-RFP, 35S::TRB2-RFP, 35S::TRB3-RFP,* and *i35S::PWO1-mCherry*, including merged images (overlay) in *Nicotiana tabacum*. The FLIM-FRET data are displayed using a lifetime Look-Up Table (LUT). Scale bar = 10 μm.

### PWO1 and TRB-enriched peaks co-localize at shared genomic targets and at *telo*-box motifs in the Arabidopsis genome

To assess the co-binding of PWO1 with TRB1-3 over its genomic targets, published ChIP-seq datasets for PWO1-FLAG (Zheng et al., 2023) and FLAG-TRBs (Wang et al. 2023) (both generated from Arabidopsis seedlings) were analyzed, and their signal profiles and peak overlaps were compared. PWO1-FLAG peaks were classified into two clusters based on TRBs binding intensity: PWO1 Cluster 1 (PWO1-TRB-enriched peaks, *n = 2,531;* Table S1) and PWO1 Cluster 2 (PWO1-biased peaks, *n = 7,067;* Table S1) (Figure 3A). Among the TRBs, FLAG-TRB1 showed the strongest enrichment across both PWO1 clusters, followed by FLAG-TRB2 and FLAG-TRB3 (Figure 3A). Genomic feature analysis revealed that PWO1 Cluster 1 peaks are predominantly located in promoters (∼85%), whereas PWO1 Cluster 2 peaks are found in promoter regions (∼62%), followed by 5’UTRs (∼16%) and Exon (∼18%) (Figure 3B). Gene Ontology (GO) Biological Process (BP) of PWO1-TRBs shared genes (Cluster 1; Table S2) highlighted ribonucleoprotein complex biogenesis, RNA splicing, and mRNA processing (Figure 3C), while PWO1-biased bound genes (Cluster 2; Table S2) were associated with stress-related pathways, including DNA damage and repair, responses to hypoxia and decrease in oxygen levels (Figure 3C). Clustering of TRB peaks based on PWO1 enrichment in cluster 1 shows that ∼23% of peaks in TRB1, ∼15% in TRB2, and ∼17% in TRB3 are enriched by PWO1, whereas TRBs cluster 2 peaks exhibits weak PWO1 binding overlap (Figure S3A). TRBs clusters peaks were associated with similar BP terms (PWO1 cluster 1; Figure 3B), except for TRB1 cluster 2, which shows a distinct functional profile (Figure S3B-D). Analysis of gene-level co-occupancy showed that, among PWO1 target genes in both clusters, 7,549 (∼82%) overlapped with TRB1, 2,722 (∼30%) with TRB2, and 2,302 (∼25%) with TRB3 (Figure 3D).

**Figure 3.**
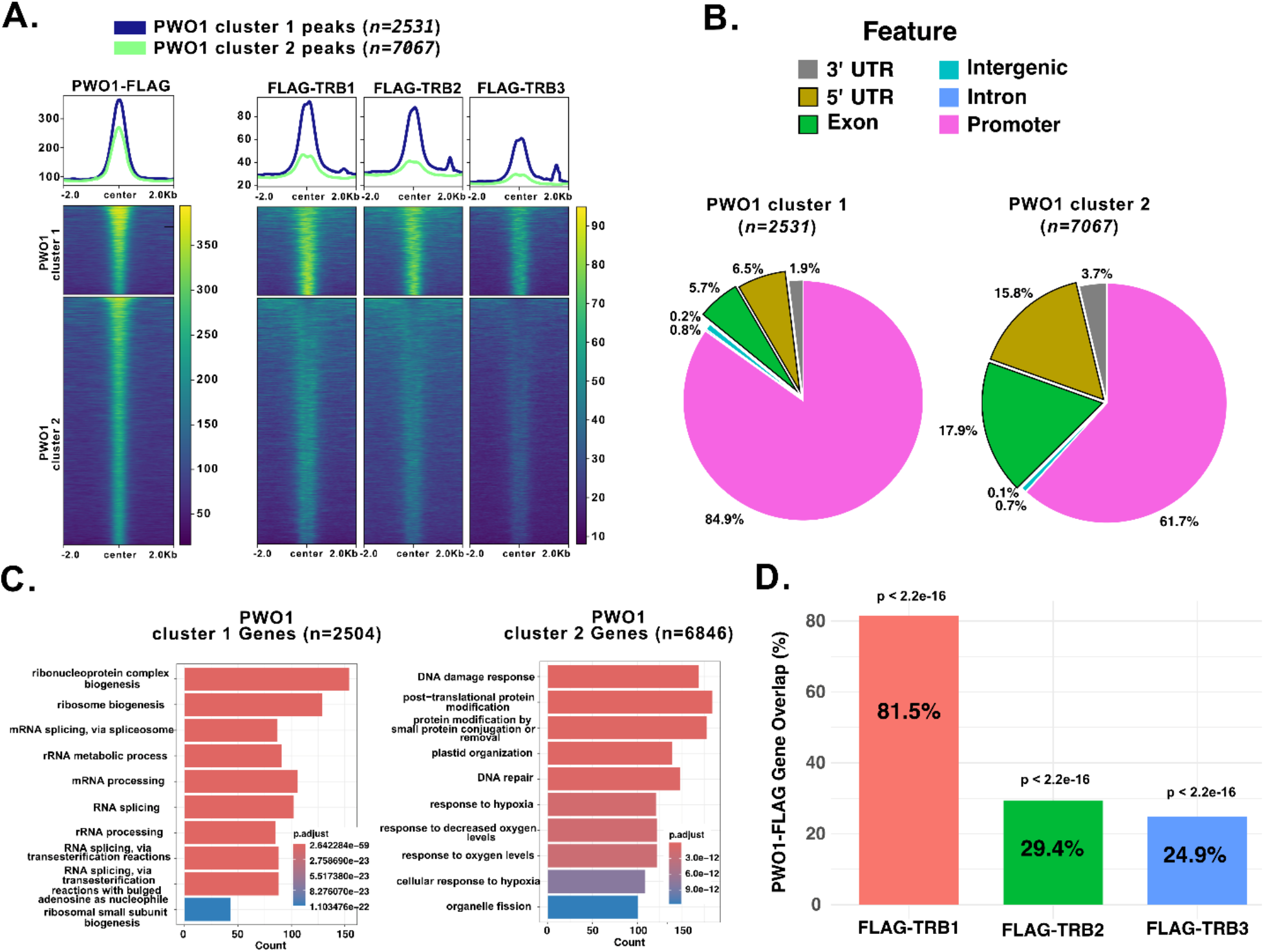
Overlapping PWO1-TRBs binding and functional annotation. Metaplots and heatmaps showing normalized ChIP-seq signals of **A.** PWO1-FLAG (Zheng *et al*., 2023), FLAG-TRB1, FLAG-TRB2, and FLAG-TRB3 (Wang *et al*., 2023) over two clusters of PWO1-FLAG peaks, classified based on TRBs enrichment over PWO1-FLAG peaks. Heatmap color scale represents normalized signal intensity (RPKM; Reads Per Kilobase of transcript per Million mapped reads), from low (blue) to high (yellow). **B.** Pie charts depicting the genomic annotation of the two clusters, PWO1 Cluster 1 and PWO1 Cluster 2, across major genomic features. **C.** GO enrichment analysis (Biological Process, BP) of genes associated with two clusters, PWO1 Cluster 1 and PWO1 Cluster 2. Bars represent the top 10 significantly enriched processes, with lengths proportional to the number of genes associated with each term. Significance was determined using clusterProfiler (Benjamini–Hochberg adjusted p < 0.05). **D.** Bar chart showing the percentage of target genes overlapping between PWO1 and TRB1-3. Hypergeometric tests (p < 2.2e-16) indicate non-random co-binding with PWO1.

Overall, these analyses indicate that PWO1 and TRB1-3 protein co-occupy a substantial fraction of regulatory regions, potentially coordinating key processes of RNA processing and translation. To explore the underlying DNA sequence features, motif enrichment analysis was performed for PWO1-FLAG (Zheng et al. 2023) and FLAG-TRB1-3 (Wang et al. 2023), revealing the *telo*-box-like motif (TAGGGTT; hereafter referred to as the *telo*-box) and the GA/TC-rich motifs among the top three enriched motifs associated with these proteins (Figure S4A-D). Motif enrichment analysis of Cluster 1 (PWO1-TRBs-bound regions) and Cluster 2 (PWO1-biased regions) revealed similar *telo-*box motif (TAGGGTT) as the second most enriched DNA motif (Figure S4E-F). A total of 13,669 *telo*-box motifs (TAGGGTT; Table S1) were identified across the TAIR10 genome (Figure S4G) and used to assess PWO1-FLAG and FLAG-TRB1-3 binding (Figure 4). Clustering based on PWO1-FLAG binding separated these *telo*-box regions into three groups: Group A (*981 telo*-box sites, 734 genes; Table S1-2) and Group B (*5,721 telo*-box sites, *3,959* genes; Table S1-2) enriched with PWO1-TRB1-3, with a stronger PWO1-FLAG signal in Group A than B; while Group C (*6,967 telo*-box sites, 1,914 genes; Table S1-2) predominantly bound only by TRB1-3 (Figure 4A). From Group A, 65.6% and 28.8% of *telo*-boxes overlap with PWO1 clusters 1 and 2, respectively. In contrast, Group B *telo*-box shows lower overlap, with 14.4% and 20.6% in PWO1 clusters 1 and 2, while Group C shows negligible overlap in either cluster. Together, these results indicate that PWO1 and TRB co-share almost half of the *telo*-boxes, while TRBs appear to associate with nearly all of them.

**Figure 4.**
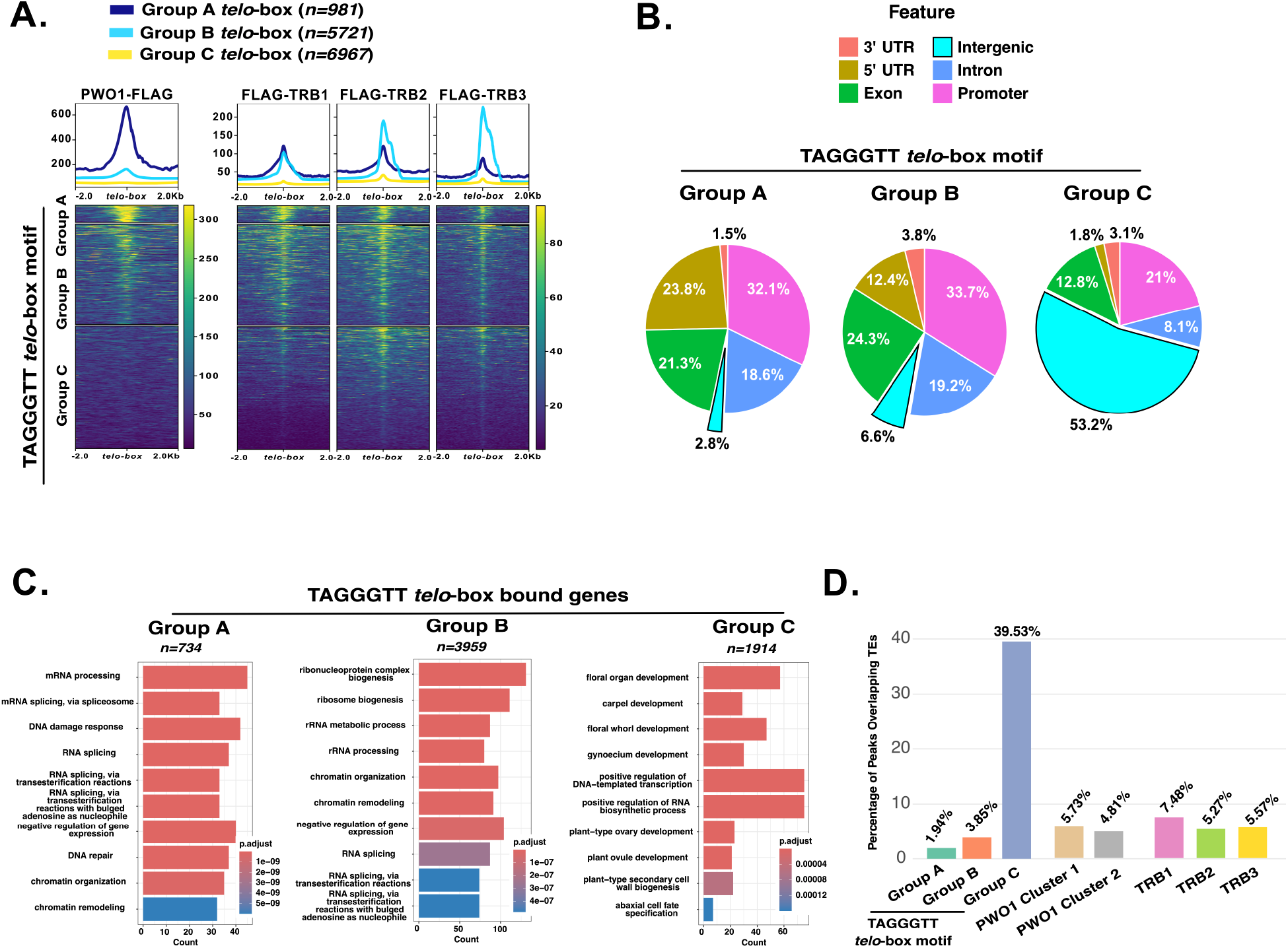
Binding of PWO1 and TRB1-3 over TAGGGTT *telo-*box motif Group A-C with associated genomic features and biological functions. **A.** Metaplots and heatmaps showing normalized ChIP-seq (Zheng *et al*., 2023; Wang *et al*., 2023) signals over Group A-C *telo-*box motif, clustered by PWO1-FLAG enrichment, illustrating the binding profiles of PWO1-FLAG, FLAG-TRB1, FLAG-TRB2, and FLAG-TRB3. Heatmap color scale represents normalized signal intensity (RPKM), from low (blue) to high (yellow) **B.** Pie charts depicting the genomic annotation of Group A-C *telo-*box motif. **C.** GO enrichment analysis of genes associated with Group A-C *telo-*box motif. Bars represent the top 10 significantly enriched biological processes, with lengths proportional to the number of genes associated with each term. Significance was determined using clusterProfiler (Benjamini–Hochberg adjusted p < 0.05). **D.** Bar plot showing the proportion of peaks from Group A-C *telo*-box motif, PWO1 cluster 1-2 and FLAG-TRB1-3 that intersect with TE annotations from the *Araport11_GTF_genes_transposons*. Numbers above bars indicate the percentage of peaks overlapping TEs.

Genomic feature analysis revealed distinct distributions for the *telo*-box motifs. Group A-B *telo*-box motifs are mainly located in gene regulatory regions, with only 2.8% and 6.6% in intronic regions (Figure 4B). In contrast, Group C *telo*-box motifs are largely found in intergenic regions (53.2%) (Figure 4B). GO-BP enrichment analysis of Group A-B *telo*-box motifs, bound by PWO1- TRBs, highlighted pathways related to mRNA processing and splicing, ribonucleoprotein complex biogenesis, DNA damage response, and chromatin organization and remodeling (Figure 4C) more similar GO-BP for PWO1-TRBs co-shared genomic regions (Figure 3C). In contrast, Group 3 *telo*-box (only TRBs-bound) motifs, were enriched for pathways related to flower development, including floral organ, whorl, gynoecium, carpel, ovule, and ovary development (Figure 4C).

Analysis of overlap with transposable elements (TEs) revealed that only a small fraction of Group A and B *telo*-box peaks intersect with TEs (2-4%), whereas Group C *telo*-box peaks show substantial TE association (∼40%). PWO1 cluster 1-2 and TRB1-3 peaks exhibited low TE overlap (∼5-7%), indicating that TE enrichment is largely specific to Group C *telo*-box motifs (Figure 4D; Table S3). Examples of Group A-B *telo*-box genes bound by PWO1 and TRB1-3 (*DE-ETIOLATED 3* (*DET3*) and *IRREGULAR XYLEM 10 (IRX10*)) are shown in Figure S5A, while an example of Group C is shown for the *SEPALLATA 3* (*SEP3*) promoter, which is bound by TRB1-3 but not by PWO1, in Figure S5B. TRB1 binding to the *telo*-box motif is primarily mediated by its MYB domain, as supported by AF3 predictions (Figure S5C) and previous studies (Mozgová et al. 2008; Wang et al. 2025). Analysis of published ChIP-seq data (Wang et al. 2025) for TRB1 variants lacking the MYB or GH1/H1-5 domains confirmed that the MYB domain is critical for global *telo*-box binding (Figure S5D), consistent with previous findings (Wang et al. 2025). Notably, TRB1-PWO1 co-bound regions (Cluster 1) were strongly affected by MYB deletion, indicating that the MYB domain is required for TRB1 association at PWO1-shared genomic sites (Figure S5E).

### PWO1-TRBs co-bound genomic targets are associated with a more active chromatin environment

To investigate the chromatin context of PWO1 and TRB1-3 co-bound regions, we examined histone mark occupancy at their peaks using published data (Chen et al. 2017; Wang et al. 2023). Two tested active histone marks (H3K4me3 and H3K27ac) were present at both PWO1 Cluster 1 (PWO1-TRBs enriched) and, more strongly, at PWO1 Cluster 2 (PWO1-biased) peaks (Figure 5A). In contrast, repressive H3K27me3 signals were largely absent from PWO1 Cluster 1 and weak at PWO1 Cluster 2 (Figure 5A). Similarly, at TRB1-3 target peaks (Figure S6A), co-bound TRBs-PWO1 regions (TRB1, TRB2, TRB3 cluster 1) are depleted of H3K27me3 and enriched for H3K4me3, whereas TRB-biased regions (TRB1, TRB2, TRB3 cluster 2) are co-occupied by both H3K27me3 and H3K4me3 (Figure S6A-B). Together, these results indicate that PWO1-TRB-enriched regions are generally depleted of repressive marks and enriched in active chromatin modifications.

**Figure 5.**
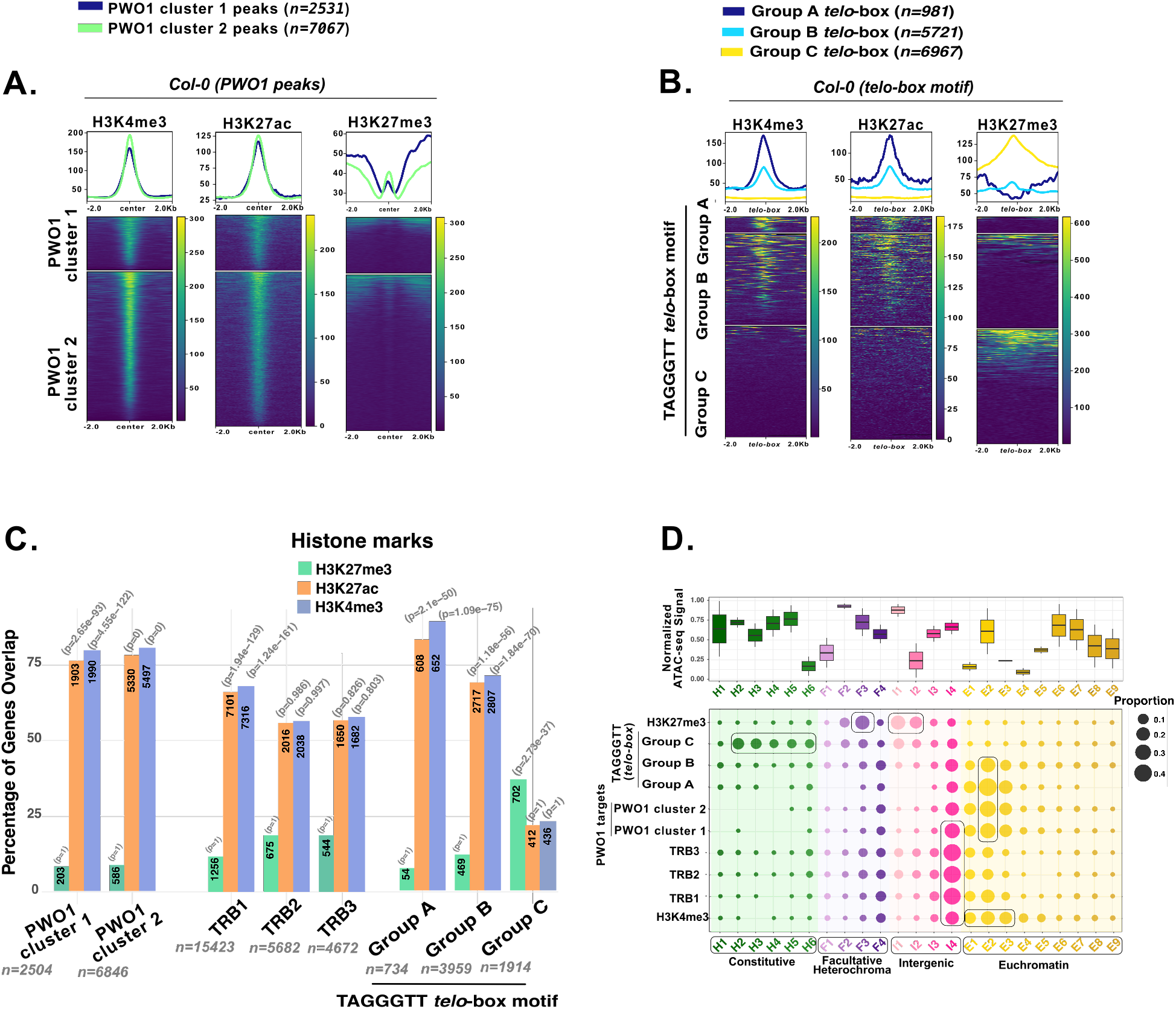
Histone mark occupancy at *telo*-box-associated regions. Metaplots and heatmaps showing normalized ChIP-seq signals (Wang *et al*., 2023; Chen *et al*., 2017) of H3K27me3, H3K4me3, and H3K27ac over **A.** PWO1-FLAG peak clusters and **B.** *telo*-box (TAGGGTT) motif clusters (Group A-C). **C.** Bar graph showing the percentage of PWO1 clusters 1-2, TRB1-3, and Group A-C *telo*-box motif–bound genes overlapping with H3K4me3, H3K27ac, and H3K27me3 occupied genes. **D.** Bubble plot showing the emission probabilities of histone modifications for PWO1 clusters 1-2, TRB1-3, Group A-C *telo*-box motifs, H3K4me3, and H3K27me3 across the 26 chromatin states of Arabidopsis. Bubble size represents emission probability (0.1–0.5), and colors indicate classification of states into major chromatin domains. Highlighted boxes denote clusters of bubbles representing genomic regions enriched in specific chromatin states. The box plot on top shows the average ATAC-seq signal for each state, representing chromatin accessibility.

We next analyzed histone mark occupancy in Groups A-C *telo*-box (TAGGGTT) motifs (Figure 5B). Group A *telo*-box motifs (i.e. bound by PWO1 (highly enriched) and TRBs; Figure 4A) were completely devoid of the repressive mark H3K27me3 and enriched in active H3K4me3 and H3K27ac marks (Figure 5B). In Group B *telo*-box motifs (i.e. bound by PWO1 (moderately enriched) and TRBs; Figure 4A) showed only ∼10.4% H3K27me3 marking (*n =593*) and were highly enriched in H3K4me3 and H3K27ac (Figure 5B; Figure S7A). In contrast, Group C *telo*-box motifs (i.e. bound only by TRBs; Figure 4A) showed ∼20.3% H3K27me3 enrichment (*n =1,412*) and lacked enrichment of H3K4me3 and H3K27ac (Figure 5B; Figure S7D). Genomic feature analysis showed that H3K27me3-marked Group B *telo*-box motifs (*n = 593*) are enriched in promoters and exons and associate with distinct GO-BP terms related to stress and plant responses to the environment (Figure S7B-C) compared with typical PWO1-TRB targets (Figure 4B-C). In contrast, Group B *telo*-box motifs (*n = 5,128*) lacking H3K27me3 resemble PWO1-TRBs shared peaks (Figure S7B-C; Figure 4B-C). Similarly, H3K27me3-marked Group C motif (*n = 1,412*) are enriched in promoter and 5’UTRs and, display GO-BP associated with floral organ development (Figure S5E-F), whereas *telo*-box motif (*n = 5,555*) lacking H3K27me3 are predominantly intergenic (∼60%) and lack GO-BP terms, consistent with their strong transposable element association (Figure S7E-F; Figure 4D).

Consistently, PWO1 peaks (clusters 1-2) and TRB1-3 targets predominantly overlap with active histone marks (H3K4me3 and H3K27ac; ∼76-80% and ∼56-68%, respectively) and show limited association with H3K27me3 (∼8–19%) (Figure 5C). Similarly, Group A and B *telo*-box–containing genes are enriched for active chromatin marks (∼69-89%), whereas Group C genes show increased association with H3K27me3 (∼37%) and correspondingly lower levels of active marks (∼21%) (Figure 5C). Chromatin state analysis (Shukla et al. 2026) further revealed that PWO1 targets and Group A-B motifs, resided in euchromatic, transcriptionally active regions associated with peaks of H3K4me3. As expected, H3K27me3-marked sites were associated with facultative heterochromatin and intergenic regions. In contrast, TRB-bound regions were frequently located at intergenic boundaries distinct from H3K27me3-bound regions. Notably, Group C motifs are predominantly associated with heterochromatic regions, known to be enriched for H3K9me2 and transposable elements (Shukla et al. 2026). In conclusion, these results indicated that PWO1-TRBs co-bound chromatin regions are predominantly associated with active chromatin; however, specific subsets coexisted with H3K27me3, highlighting a context-dependent chromatin environment at their target sites.

### PWO1 binding depends on TRBs at shared genomic regions

To assess a TRB-dependent recruitment of PWO1 to chromatin and *telo*-box-associated regions, *PWO1pro::PWO1-GFP* lines were generated in WT, *trb1-2 trb2-2 trb3* triple mutant, and *trb1-2 trb2-2 trb3^+/−^* backgrounds (Figure S8A-C). PWO1-GFP showed stable nuclear localization in root tips across all genotypes (Figure S8D). ChIP-seq analysis of PWO1-GFP was performed in WT and *trb1 trb2 trb3* mutant backgrounds (a pool of *PWO1pro::PWO1-GFP/trb1-2 trb2-2 trb3-*1 and *PWO1pro::PWO1-GFP/trb1-2 trb2-2 trb3-1^+/−^;* hereafter referred to as the *trb123* mutant background). Two biological replicates showed high reproducibility, as evidenced by principal component analysis (PCA) clustering and strong pairwise correlations (Figure S9A-B). Interestingly, in the *trb123* mutant background, PWO1 showed a substantial loss of binding to its targets, with only 58 targets identified compared to 1,739 in WT (Figure 6A; Table 4). This loss of PWO1 binding in the *trb123* background predominantly affects genes associated with BP related to chromatin organization/remodeling and DNA damage response (Figure S9C). This reduced binding of PWO1-GFP is further supported by decreased enrichment levels in the *trb123* background, particularly across gene bodies relative to the TSS peaks observed in WT, indicating that TRBs are required for efficient PWO1 recruitment to chromatin (Figure 6B-C). Notably, the strongest impact on PWO1 binding was observed at loci co-occupied by PWO1 and TRBs over PWO1-FLAG targets (Zheng et al. 2023) (Figure 6D). A similar trend was evident at *telo*-box-containing regions, where PWO1-GFP enrichment was preferentially lost at group A *telo*-boxes co-bound by PWO1 and TRBs in *trb123* background (Figure 6E-F), indicating that TRBs facilitate PWO1 targeting to these short motifs. Lastly, we compared the PWO1 target genes identified from our PWO1-GFP dataset with previously published PWO1 binding datasets, including PWO1-FLAG (Zheng et al. 2023) and PWO1-GFP (Yang et al. 2024). Among the PWO1 target genes identified in our dataset, 1,415 genes (∼83%) were also reported in previous studies (Figure S9D), indicating considerable overlap between datasets despite differences in experimental approaches and the number of identified targets.

**Figure 6.**
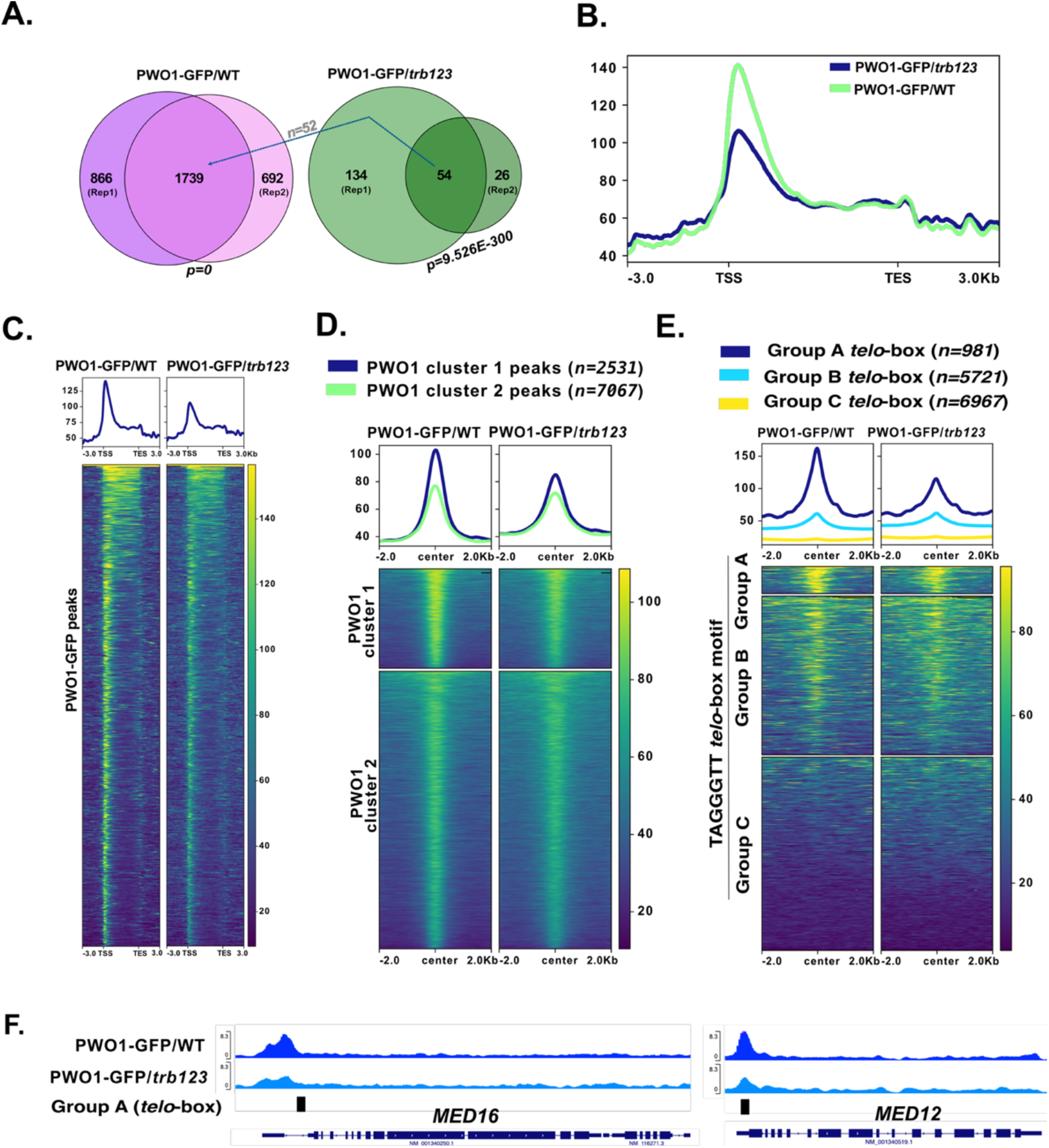
PWO1-GFP chromatin binding in the *trb123* mutant background. **A.** Venn diagram showing the overlap of PWO1pro::PWO1-GFP target regions between two biological replicates in WT and *trb123* backgrounds (a pool of *PWO1pro::PWO1-GFP/trb1-2 trb2-2 trb3-1* and *PWO1pro::PWO1-GFP/trb1-2 trb2-2 trb3-1^+/−^*; referred to as the *trb123* mutant background). Out of 54 targets identified in the *PWO1-GFP/trb123* condition, 52 overlaps with PWO1-GFP/WT targets. **B-C.** Metaplots and heatmaps showing normalized PWO1-GFP ChIP-seq signals over PWO1 target regions (*n = 1739*). **D-E.** Enrichment of PWO1-GFP over **D.** PWO1-FLAG targets (Zheng *et al*., 2023) and **E.** *telo*-box motif in WT and *trb123* backgrounds. **F.** IGV browser snapshots showing PWO1-GFP binding at the MED16 and MED12 loci in WT and *trb123* mutant backgrounds.

### PWO1 and TRBs cooperatively regulate multiple aspects of plant development, including ectopic lignification

To assess the coordinated role of PWO1 and TRBs in Arabidopsis development, we generated double mutants (*pwo1-1 trb1-1, pwo1-1 trb2-1, pwo1-1 trb3-1*) (Figure S10A-B) and compared their phenotypes with single mutants and wild type (*Col-0*) plants (Figure S11A-F). The *pwo1-1* mutant showed early bolting and flowering, reduced leaf number, and shorter roots (Figure S11A-B, D-E) as shown earlier (Hohenstatt et al. 2018), whereas *trb2-2* displayed slightly delayed bolting and flowering under our growth conditions (Figure S11A-B). Other *trb* single mutants resembled wild type (Figure S11A-E), except *trb1-1*, which had fewer leaves (Figure S11D). Double mutants largely mirrored *pwo1-1*, with the exception of further leaf reduction in *pwo1-1 trb1-1* and partial rescue of root length (Figure S11A-E). These results indicate that TRBs are functionally redundant in regulating these developmental traits.

We next analyzed the developmental phenotypes of higher-order *pwo1-1 trb1-2 trb3-1* triple mutants (Figure 7, Figure S10C-D). Root length measurements of 14-day-old seedlings revealed that *pwo1-1 trb1-2 trb3-1* triple mutants developed shorter and more variable roots compared with Col-0 plants (Figure 7A-B). Strikingly, the pwo*1-1 trb1-2 trb3-1* triple mutants exhibited a significant reduction in rosette area, petiole length and leaf numbers compared with control plants (Col-0*, pwo1-1*, *trb1-2 trb3-1*) (Figure 7C-E, Figure S12A). No phenotypic differences were observed in *pwo1-1 trb1-2 trb3-1* triple mutants for bolting and flowering compared to *Col-0* (Figure 7F-H). One of the most prominent post-bolting phenotypes observed in the *pwo1-1 trb1-2 trb3-1* triple mutant was arrest of main stem internode elongation, resulting in overall stunted growth (Figure 7I-J). Approximately 20% of these triple mutant plants exhibited severe shortening of the main stem, whereas 40% displayed severely reduced stem elongation accompanied by loss of apical dominance and excessive axillary branching. The remaining 40% of plants developed a shorter but morphologically normal main inflorescence stem (Figure S12B). Furthermore, siliques of the *pwo1-1 trb1-2 trb3-1* triple mutants on the main stem were fewer and morphologically aberrant - they were flattened, swollen, and bent at tips (Figure S12C). Overall, these results indicate that PWO1, together with at least TRB1 and TRB3, plays a major, partially redundant role in stem elongation, axillary branching, silique morphology, and overall plant development. The phenotypical differences are likely explained by different timing of meristem arrest or perturbation. Interestingly, in the *pwo1-1 trb1-1/trb1-2 trb3-1* triple mutant allele background, segregating *trb1-2^+/−^* and *trb3-1^+/−^* alleles exhibit distinct phenotypes. The *pwo1-1 trb1-2^+/−^ trb3-1* plants showed a wild-type-like phenotype, whereas *pwo1-1 trb1-1 trb3-1^+/−^* plants resembled the *pwo1-1 trb1-2 trb3-1* and *pwo1-1 trb1-1 trb3-1* triple mutants (Figure S12D). These observations indicate that full TRB3 dosage is required for proper development in the *pwo1-1 trb1-1* mutant background.

**Figure 7.**
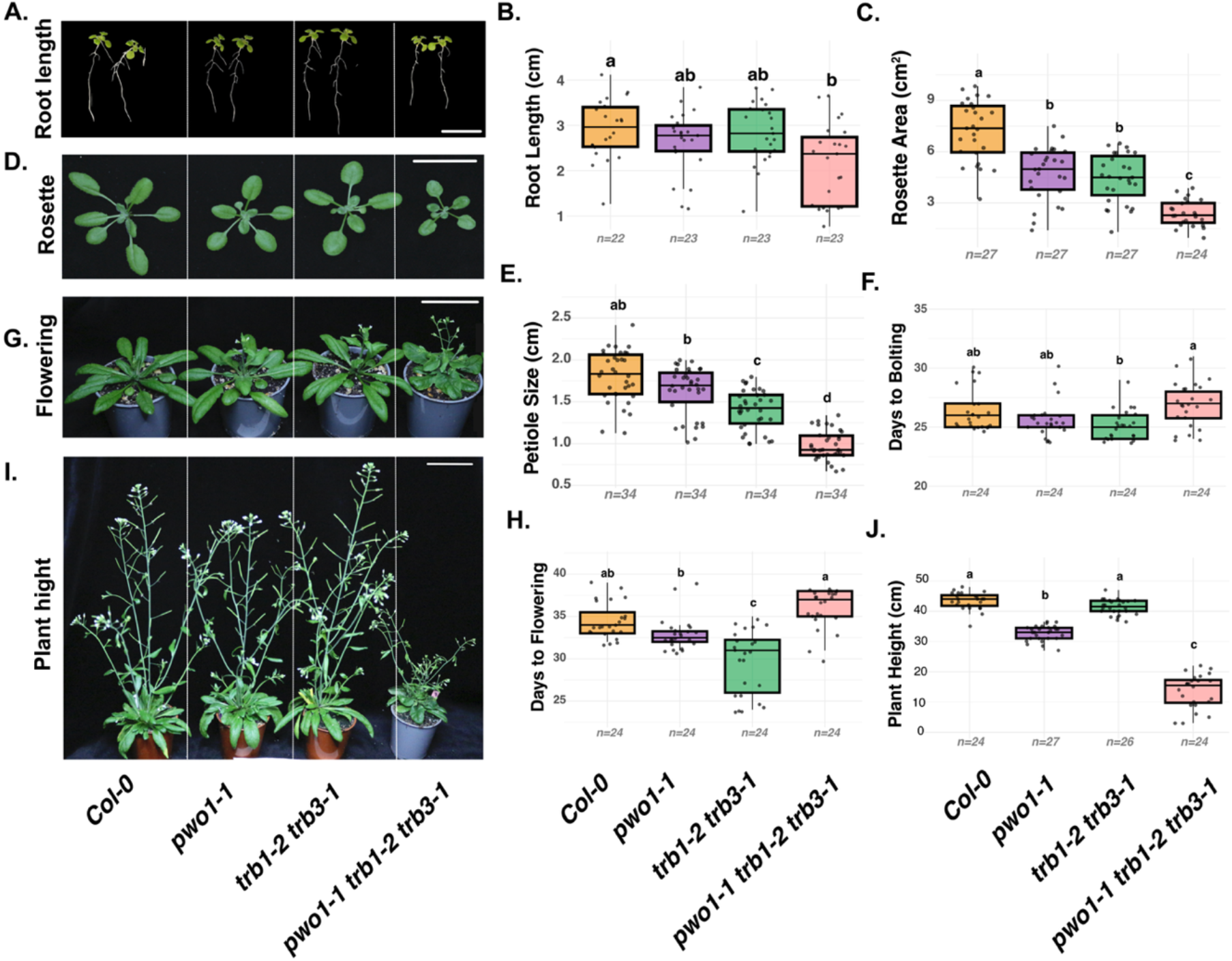
Phenotypic analysis of *pwo1-1 trb1-2 trb3-1* triple mutants in *Arabidopsis* plants. **A, D, G,I**. Representative images of **A.** root length (14 DAG), **D.** rosette area (20 DAG), **G.** flowering time (34 DAG) and **I.** plant height (44 DAG), phenotypes for Col-0, *pwo1-1*, *trb1-2 trb3-1*, and *pwo1-1 trb1-2 trb3-1* plants. **B**, **C**, **E**, **F**, **H**, **J**. Boxplots showing quantitative analyses of **B.** root length **C.** rosette size, **E.** petiole length, **F.** days to bolting, **H.** days to flowering and **J.** plant height of Col-0, *pwo1-1*, *trb1-2 trb3-1*, and *pwo1-1 trb1-2 trb3-1* plants. Black points represent individual plants and are displayed with horizontal jitter. Box colors indicate genotype groups: orange for Col-0, purple for single mutants (*pwo1-1*), green for double mutant (*trb1-2 trb3-1*) and pink for triple mutant (*pwo1-1 trb1-2 trb3-1*). Different letters above the boxes indicate statistically significant differences between genotypes (one-way ANOVA followed by Tukey’s HSD test, *p* < 0.01). Scale bar = 5 cm. DAG (Days After Germination)

The stunted or dwarf phenotypes can arise from premature or ectopic lignification (Caño-Delgado et al. 2000; Perkins et al. 2020). To assess whether altered lignin deposition contributes to the stunted growth of the *pwo1-1 trb1-2 trb3-1* triple mutant, transverse stem sections across developmental stages were stained with Basic Fuchsin (Figure 8). At 28 days, shortly after bolting, the triple mutant already displayed premature lignin accumulation in interfascicular fibers (IF) compared to Col-0*, pwo1-1*, *trb1-2 trb3-1* (Figure 8A; Figure S13A). By 34 days after germination, lignification expanded markedly, showing strong and continuous deposition in both IF and phloem (Figure 8B, 8E; Figure S13A). This ectopic and enhanced lignification exacerbated through intermediate and mature stages (40 DAG, 48 DAG and 58 DAG) (Figure 8C-E; Figure S13A-B).

**Figure 8.**
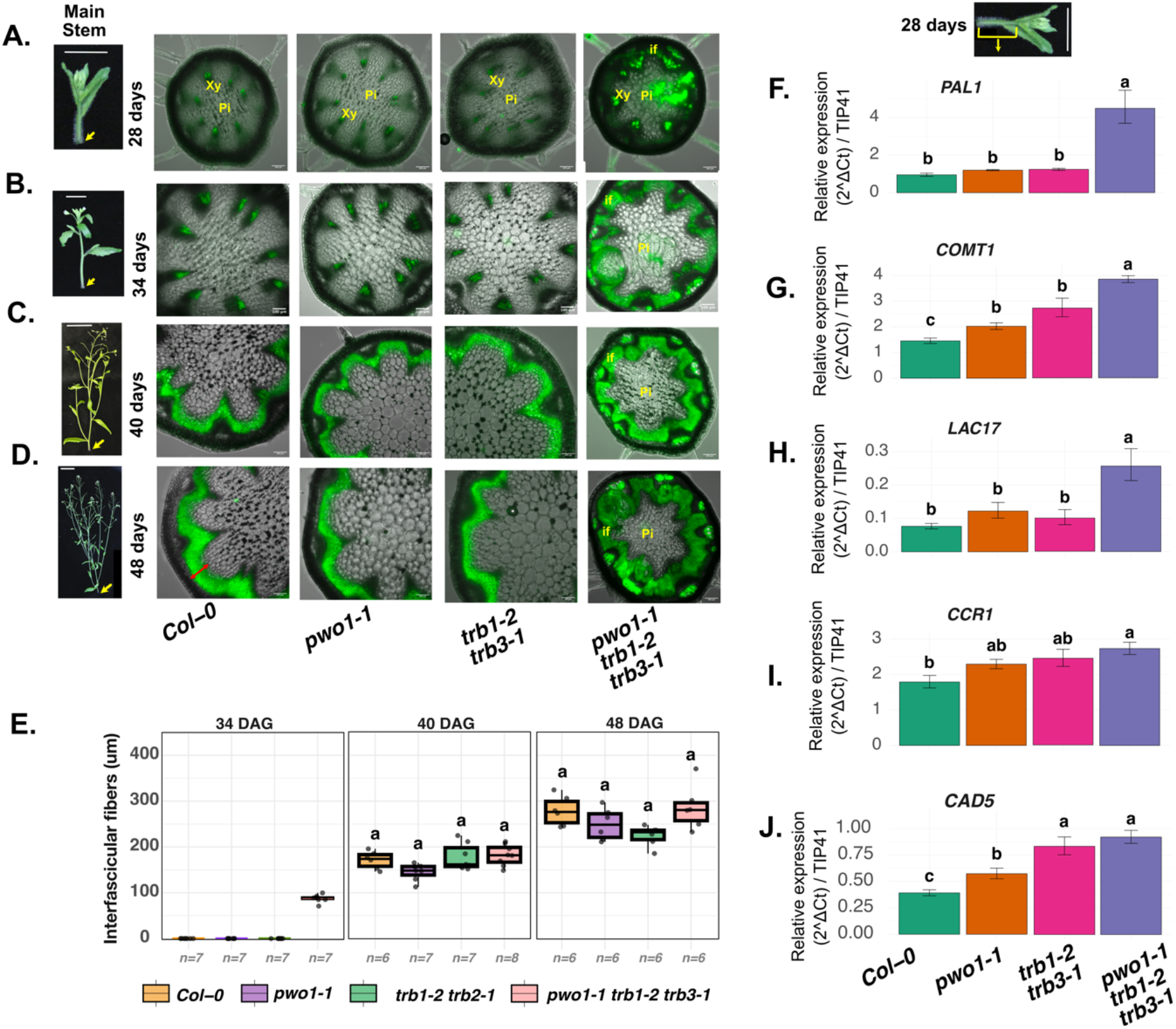
Temporal analysis of interfascicular fiber lignification in the main stem. Transverse sections of the basal region of the main stem stained with Basic Fuchsin, which labels lignified interfascicular fibers (IF), are shown for *pwo1 trb1 trb3* triple mutants, *pwo1*, *trb1 trb3*, and Col-0 at **A.** 28 days after sowing (prior to flowering), **B.** 34 days, **C.** 40 days, and **D.** 48 days. Yellow arrows indicate the positions along the stem from which transverse sections were taken. **E.** Quantification of Basic Fuchsin–stained interfascicular fiber area (red arrow) across all developmental time points. **F.** qPCR analysis of lignin biosynthesis–related gene expression in 28-day-old stems collected after bolting but prior to flowering. Xy = xylem; Pi = pith. DAG (Days After Germination)

To determine whether this phenotype reflects transcriptional misregulation of the lignin biosynthetic pathway (Figure S14I), gene expression was analyzed at 28 days stem after bolting before flowering, coinciding with the onset of lignin deposition (Figure 8A). *PHENYLALANINE AMMONIA-LYASE 1* (*PAL1*), *CAFFEIC ACID O-METHYLTRANSFERASE 1* (*COMT1*), and *LACCASE 17* (*LAC17*) were significantly upregulated in the *pwo1-1 trb1-2 trb3-1* triple mutant, whereas *CINNAMOYL-CoA REDUCTASE 1* (*CCR1*) and *CINNAMYL ALCOHOL DEHYDROGENASE 5* (*CAD5*) showed increase expression comparable to *pwo1-1* and *trb1-2 trb3-1* (Figure 8F-J). No changes in expression of *PAL1*, *COMT1*, and *LAC17* were observed at 14 DAG (Figure S13A-C), indicating stage-specific upregulation. Upstream regulators of secondary cell wall biosynthesis genes, such as *NAC SECONDARY WALL THICKENING PROMOTING FACTOR 1* (*NST1*) and *VASCULAR-RELATED NAC-DOMAIN 7* (VND7), along with their downstream target *MYB DOMAIN PROTEIN 83* (*MYB83*), were also upregulated in the *pwo1-1 trb1-2 trb3-1* triple mutant (Figure S13F-H) in 28 DAG stems, whereas these genes were unaffected in 14 DAG stems (Figure S13D-E). This coordinated upregulation across regulatory, intermediate, and structural components of the lignin biosynthesis pathway in *pwo1-1 trb1-2 trb3-1* triple mutant is consistent with premature activation of the secondary cell wall program (Figure S13I). Together, these results indicate that the *pwo1-1 trb1-2 trb3-1* mutant undergoes premature activation of the secondary cell wall regulatory network, leading to early and ectopic lignin deposition, which likely underlies the reduced internode elongation and stunted growth phenotype.

## Discussion

### PWO1 forms conserved telomeric complexes with TRBs

In complex epigenetic regulatory networks, accessory proteins play a crucial role in targeting chromatin-modifying factors to specific loci, thereby controlling transcriptional outcomes (Scheid et al. 2021; Godwin and Farrona 2022; Khan et al. 2025). In this study, we have extensively examined the molecular roles of two such proteins namely, PWO1 and TRBs, which have recently been shown to participate in distinct epigenetic complexes (Zheng et al., 2023; Wang et al., 2025; Wang et al., 2023; Zhou et al., 2018; Hohenstatt et al., 2018). Despite their involvement in multiple complexes, the coordinated functions of PWO1 and TRBs remain poorly understood. Here, we observed that the PWO1-TRB interaction is conserved since lycophytes (Figure S1A), complementing our earlier report of PWO1-PRC2 interaction in the same lineage (Khan et al. 2026). Notably, PWO1 first emerges in lycophytes, while other PEAT-related components, including TRBs, ARID domain-containing proteins and UBP5 are already conserved in this lineage (Kusová et al., 2023; Amiard et al., 2024; Zheng et al., 2023; Khan, Ghosh, et al., 2025; Khan, Haider, et al., 2025). This raises the possibility that a PEAT-like complex incorporating PWO, TRB, ARID, UBP5 and other PEAT factors could have formed early in lycophyte evolution, contributing to the establishment of conserved chromatin-regulatory networks and specific developmental adaptations during early plant terrestrialization.

Recently, TRBs from the bryophyte *Physcomitrium patens* were shown to localize in *N. benthamiana* in a pattern similar to that of Arabidopsis TRBs (Kusová et al. 2025). In our study, one TRB paralog from *S. moellendorffii* exhibited a localization pattern similar to that observed in angiosperms, suggesting a conserved subnuclear organization across land plants. It formed speckle-like structures within the nucleoplasm and showed strong nucleolar accumulation when transiently expressed in *N. benthamiana* (Figure S2A-E). Although our analysis included only a single *S. moellendorffii* TRB paralog (Figure 1; Figure S1-2) and therefore does not capture the full spectrum of SmPWOa-SmTRB interactions, these findings nonetheless provide important evidence supporting the existence of PWO-TRB interactions in early-diverging lycophyte lineages. TRB1-derived nuclear speckles were previously reported to overlap with telomeric regions (Procházková Schrumpfová et al. 2014; Khosravi et al. 2020), supporting the idea that TRBs function as conserved telomere-binding proteins in plants. Notably, PWO1 and lycophyte PWOs also form nuclear speckles (Hohenstatt et al. 2018; Khan et al. 2025) like those seen for TRBs (Procházková Schrumpfová et al. 2014), although the nature and positioning of PWO1 speckles within the nucleus were not previously understood. Our co-localization experiments with a telomere marker show that PWO1 and TRBs occupy the same subnuclear space, forming overlapping nuclear speckles (Figure 2), implying that, like TRBs, PWOs also aggregate at plant telomeres. Furthermore, the interaction between PWO1 and TRBs appears to occur specifically at speckles (Figure 1H, 1I), which correspond to telomeric regions (Khosravi et al. 2020). These findings position PWOs as one of the closest TRB-like proteins in terms of telomeric association, and, to our knowledge, no other TRB-interacting protein reported so far exhibits a comparable telomere-associated localization pattern. This provides compelling evolutionary evidence, showing that despite major divergence in nuclear organization and telomeric structure across plant species, PWO1 and TRBs consistently form speckle-like structures associated with telomeric regions (Figure1-2). This suggests that, despite millions of years of evolutionary divergence, these proteins may recognize conserved higher-order chromatin features such as telomeres across land plants, pointing to a preserved role in nuclear organization that requires further investigation. One discrepancy between Y2H and FLIM-FRET results concerns PWO1 interactions with TRBs. In Y2H, PWO1 interacted with TRB1 and TRB2 but not TRB3 (Figure S1A) as shown earlier (Tan et al., 2018; Zheng et al., 2023), whereas in *planta* FLIM-FRET revealed an interaction with TRB3 as well, suggesting that PWO1 can potentially interact with all three TRBs paralogs (TRB1-3) (Figure 1H). The lack of PWO1-TRB3 interaction in Y2H may reflect misfolding, artifacts of the yeast heterologous system, or the absence of plant-specific factors required to stabilize this interaction. Furthermore, compared to the Y2H assay, FLIM/FRET is able to detect even very weak, but biologically highly relevant interactions in the protein native conditions (Yamoune et al. 2024; Cairo et al. 2026; Khan et al. 2026). Additionally, PWO1 was shown to interact with TRB4 and TRB5 paralogs (Kusová et al., 2023), highlighting a close association between PWO1 and the TRB protein family in Arabidopsis. Taken together, these data (Figure 1-2; Figure S1-2) indicate that PWO is capable of physically interacting with TRB proteins across plant evolution. This interaction is also conserved within subnuclear space specifically at telomeric regions.

### PWO1 and TRB1-3 co-occupy active chromatin regions and are essential for normal plant development

Our findings indicate that TRB proteins broadly define *telo*-box occupancy across the genome (Procházková Schrumpfová et al. 2014; Schrumpfová et al. 2016), while PWO1 associates with a selective subset of these sites (Figure 4). TRB1-3 bind nearly all *telo*-box motifs genome-wide. In contrast, only approximately half of the *telo*-boxes are co-occupied by PWO1, indicating the possibility that PWO1 may associate with or redirect TRBs to a specific set of *telo*-box motifs (Figure 4A). PWO1-TRBs co-bound *telo*-box motifs are enriched in gene-proximal, transcriptionally regulated regions and associated with genes involved in RNA biogenesis and ribonucleoprotein function. By contrast, TRB-only bound *telo*-box regions tend to reside in compact, intergenic chromatin, including more TE-associated sites, and are linked to genes involved in developmental regulations (Figure 4B-D). This observed pattern makes it tempting to speculate that PWO1 modulates TRBs placement towards transcriptionally regulated domains, thereby diversifying their functional output.

Analysis of PWO1 binding peaks further supports the idea that PWO1 association specifies the function of TRBs at a subset of co-occupied chromatin loci. Approximately 23% of PWO1-bound regions are co-enriched with TRBs, whereas the remaining sites display a more PWO1-biased profile with reduced TRB occupancy (Figure 3A). A similar pattern was observed for TRB1-3 peaks, where PWO1 is enriched at only ∼15–20% of TRB sites, while the remainder are more TRB-biased (Figure S3A). PWO1-TRB co-enriched peaks among PWO1 targets are associated with genes involved in RNA processing and biogenesis. In contrast, PWO1-dominant peaks with limited TRB occupancy are linked to stress-responsive pathways, including hypoxia (Figure 3B-D). A similar trend is observed for TRB1 peaks, which show distinct biological functions depending on whether they are co-occupied by PWO1 or not (Figure S3B). These patterns suggest that PWO1 and TRBs engage in functionally distinct regulatory modes: when co-occupied, they direct gene regulation toward core RNA metabolic processes, whereas independently they have different roles (Figure 3C, Figure S3B). Although the MYB domain of TRB1 is critical for high-affinity binding to *telo*-box motifs (TAGGGTT), as its deletion markedly reduces TRB1 occupancy at these sites (Figure S5D), this does not exclude the possibility that other proteins, such as PWO1, may contribute to define their binding preference. PWO1-TRB1 co-bound genomic regions are particularly sensitive to MYB deletion (Figure S5E), suggesting that MYB domain is required for PWO binding.

One of the key questions addressed in our analysis concerns the dependency of PWO1 binding on TRBs across chromatin. Interestingly, the presence of TRBs appears to be critical for PWO1 binding at a subset of chromatin regions (Figure 6A-C), consistent with the notion that TRBs may function as typical transcription factors that recruit additional cofactors to specific loci. Indeed, our analysis shows that deletion of *trb1/2/3* results in a substantial loss of PWO1 binding at co-shared regions (Figure 6D-E), including both *telo*-box–associated sites (Figure 6F-G). Interestingly, these co-shared regions account for only ∼15-20% of total PWO1 peaks (Figure 3). Thus, the majority of PWO1 binding occurs independently of detectable TRB occupancy, raising the question of how PWO1 is recruited to chromatin in these contexts. We speculate that PWO1 may associate with modified histone tails via its PWWP domain at these regions (Hohenstatt et al. 2018; Zheng et al. 2023) or with transcription factors that bind GA-rich motifs, such as Basic Pentacysteine proteins (BPCs) (Mu et al. 2017), potentially functioning independently of TRBs. However, this possibility requires further experimental validation. The lower number of PWO1 genomic targets detected in our PWO1-GFP ChIP (Figure 6A) compared with previous PWO1-GFP (Yang et al. 2024), performed in a *pwo1* mutant background, and PWO1-FLAG (Zheng et al. 2023) datasets may result from differences in the experimental system and tag properties. Since our construct is expressed in the native PWO1 background, endogenous protein may compete for target binding and reduce detection of tagged PWO1-GFP enrichment (Figure S9D). In addition, differences in tag size and antibody lot-to-lot variation (e.g., FLAG versus GFP) may affect ChIP efficiency and the number of detected genomic targets.

Chromatin profiling of PWO1-TRB1-3 co-occupied regions highlights their functional diversification. The co-bound PWO1 peaks are enriched for active histone marks such as H3K4me3 and H3K27ac and are depleted of the repressive H3K27me3 mark (Figure 5A). A similar trend is observed at TRB1-3 peaks: regions co-shared with PWO1 are depleted of H3K27me3, whereas TRB1-3 peaks with little or no PWO1 binding are enriched for both H3K27me3 and H3K4me3, suggesting more dynamic chromatin states (Figure S6). Although TRBs have been reported to recruit PRC2 and mediate H3K27me3 deposition at specific loci (Zhou et al. 2018; Wang et al. 2023), PWO1-TRB co-bound regions in our study show reduced H3K27me3 levels. This suggests that PWO1 association may promote a less repressive chromatin environment at TRB-bound regions. In contrast, PWO1-biased regions, while still enriched in active histone modifications, retain a modest level of H3K27me3 (Figure 5A), indicating a more heterogeneous chromatin context. A similar distinction is evident when *telo*-box clusters are examined: sites co-bound by PWO1 and TRBs predominantly localize to chromatin regions depleted of H3K27me3 and enriched in active marks, whereas TRB-only *telo*-box regions are associated with H3K27me3-rich, transcriptionally less active chromatin (Figure 5B, D). Interestingly, a small subset of *telo*-box motifs (n = 593) is co-occupied by PWO1 and TRBs and enriched for PRC2-associated H3K27me3 marks, linking them to cutin biosynthesis, stress responses, and insect defense (Figure S7A). These loci provide an ideal context to explore how PWO1, TRBs, and PRC2 function together, as the implications of this interaction remain largely unknown. Consistent with this observation, the *SEP3* promoter, which has previously been shown to recruit PRC2 to maintain H3K27me3-mediated repression (Zhou et al. 2018), contains a *telo*-box that falls into a category not co-bound by PWO1, thereby potentially permitting TRB-mediated PRC2 recruitment and H3K27me3 deposition (Figure S5B). Together, these observations are consistent with previous studies showing that PWO1 and TRBs participate in chromatin-modifying complexes, such as PEAT-UBP5 (Zheng et al., 2023), and integrate opposing chromatin modification pathways (Hohenstatt et al. 2018; Zhou et al. 2018; Wang et al. 2023) to regulate gene expression. Our results further suggest that PWO1 association with TRBs is linked to distinct chromatin environments compared with regions occupied by TRBs alone, providing additional regulatory specificity. As PWO1 and TRB-3 associate with thousands of genomic loci (Figure 3), their functions are likely dynamically regulated during development and stress responses. However, existing ChIP-seq datasets, largely generated from young seedlings (∼2–3 weeks), may not fully capture developmental or cell type-specific patterns of PWO1-TRB occupancy. Future stage- and cell type-resolved profiling will therefore be important to define their regulatory landscape.

Our genetic analyses indicate that PWO1 and TRBs act closely to regulate Arabidopsis development, with PWO1 exerting dominant effects on flowering, leaf number, and root growth, while TRBs modulate specific traits in the *pwo1 trb* double mutant background (Figure S11). Consistent with this, higher-order mutants (*pwo1-1 trb1-2 trb3-1*) exhibit severe developmental defects, including stunted growth due to main stem internode arrest, increased axillary branching, and aberrant silique morphology (Figure 7; Figure S12), together with premature and ectopic lignin deposition (Figure 8), likely reflecting altered timing of cellular differentiation. Importantly, the stunted growth phenotype may represent an indirect consequence of post-bolting developmental constraints triggered by early lignification soon after bolting (Figure 8A; S13A), which subsequently impacts later developmental outcomes in the triple mutant. Furthermore, analysis of *trb1* and *trb3* mutant combinations reveals a combinatorial dosage effect, where reducing TRB3 dosage in the *pwo1-1 trb1-1 trb3-1^+/-^* background, due to one disrupted TRB3 allele, leads to strong developmental defects, whereas TRB1 heterozygosity alone does not (Figure S12D). These findings indicate that TRB1 and TRB3 contribute to a context-dependent and dosage-sensitive manner. Interestingly, recent reports have revealed complementary functional biases among TRB paralogs, with TRB1 being more strongly associated with PEAT-related functions and NuA4-mediated gene regulation, whereas TRB3 is more closely linked to PRC2-associated chromatin regulation (Mendler et al. 2026). This supports the relevance of the *pwo1 trb1 trb3* triple mutant, in which the observed phenotype may arise from defects in both pathways, making it particularly informative for dissecting the opposing chromatin regulatory mechanisms mediated by PWO1. Together, these observations suggest that PWO1-TRB1-3 function is not limited to a single developmental pathway but reflects a broader chromatin-associated regulatory module whose output depends on developmental stage and tissue context. This is consistent with previous reports showing that combined loss of *TRB1* and *TRB3* activities (i.e. *trb1 trb3*) in *lhp1* mutant background enhances developmental phenotypes (Zhou et al. 2016), further supporting a cooperative role of these factors in plant development.

The upregulation of lignin biosynthetic genes (*PAL1*, *COMT1*, and *LAC17*) along with secondary cell wall regulators (*NST1*, *VND7*, and *MYB83*) in the *pwo1 trb1 trb3* triple mutant (Figure 8F-H; Figure S13F-H) suggests premature activation of the secondary cell wall transcriptional network. NST1, a master regulator that functions redundantly with NST2 and NST3, is known to induce ectopic lignification and restrict growth when overexpressed, while combined loss-of-function mutants show reduced lignification in IFs (Mitsuda et al. 2005, 2007; Zhong et al. 2007; Liu et al. 2021). During normal development, *NST1* expression is particularly enriched in IFs and has also been observed in anthers and siliques (Mitsuda et al. 2005). The lignification apparent in the pith of *pwo1-1 trb1-2 trb3-1* early after bolting implies strong developmental aberrations, possibly associating with changes in the cell identity and/or activation of NAC master switches, triggering the downstream transcriptional cascade and ectopic secondary cell wall formation. We further postulate that ectopic or mis-regulation of *NST1* expression in leaves and reproductive organs in the *pwo1 trb1 trb3* triple mutant could contribute to the altered leaf morphology and silique bending phenotypes (Figure S12A, S12C), although this will need to be tested. These findings suggest that PWO1 and TRBs cooperatively maintain proper timing of secondary cell wall activation, and their absence disrupts developmental regulation, leading to reduced internode elongation and the stunted phenotype.

Overall, our results reveal that PWO1 and TRBs cooperate to shape chromatin landscapes and developmental outcomes in Arabidopsis. PWO1 appears to modulate TRB targeting to specific genomic and *telo*-box loci, favoring transcriptionally active chromatin states and influencing gene expression programs. The severe phenotype of the *pwo1-1 trb1-2 trb3-1* triple mutant underscores the functional importance of this cooperation, while dosage-sensitive effects of TRB1 and TRB3 highlight the need for precise complex stoichiometry. Together, these findings point to a synergistic PWO1-TRB module linking epigenetic regulation with development, with TRBs serving as DNA-connecting hub involved in both gene activation and repression, depending on the complex partners. How differential and dynamic association of different chromatin proteins with TRBs is regulated, remains an important question for future studies.

## Methodology

### Plant Materials and Growth Conditions

The *Arabidopsis thaliana* mutants *pwo1* (SAIL_342_C09), *trb1-1* (SALK_001540), *trb1-2* (SALK_001540), *trb2-2* (SALKseq_4604), *trb3-1* (SALK_134641; SALK_134641.43.00), and *trb3-2* (SALK_134645) were obtained from the SALK T-DNA insertion line collection in the *Columbia-0* (*Col-0*) background (Alonso et al. 2003; Hohenstatt et al. 2018; Zhou et al. 2018). The FLAG mutant lines (*trb2-1*; FLAG_242F11 and trb2-2; FLAG_242F11; CS882628) in the Wassilewskija-2 (Ws-2) background were obtained from the INRA T-DNA insertion line collection (Samson 2002). Single mutant lines (*pwo1-1*, *trb1-1*, *trb2-2*, and *trb3-1*) were crossed to generate double mutants (*pwo1 trb’s* double mutant *combinations*). Subsequently, *pwo1-1* was crossed with *trb1-1 trb3-1* and *trb1-2 trb3-1* double mutants to generate *pwo1 trb1 trb3* triple mutants. The *PWO1*pro::*PWO1-GFP* line (Hohenstatt et al., 2018) was crossed with the *trb123* triple mutant (Wang et al. 2023) to generate the *PWO1*pro::*PWO1-GFP/trb123* line. Genotyping of mutants were performed using oligonucleotide primers listed in Table S5. Seeds were surface sterilized in 70% ethanol for 5 minutes, followed by 100% ethanol for 1 minute, air-dried, and sown on half-strength Murashige and Skoog (½ MS) medium supplemented with 0.8% agar and 0.5% sucrose. To break dormancy and synchronize germination, seeds were stratified at 4 °C for 3 days in darkness. For root assays and qRT-PCR analysis, seedlings were grown in a controlled growth chamber under long-day conditions (16 h light / 8 h dark). For phenotypic analysis, stratified seeds were sown on soil and transferred to long-day conditions. Rosette diameter was measured in 20-24-day-old plants, followed by monitoring bolting and overall plant growth. Flowering time was determined by recording the number of days until the first flower opened, and main shoot length and plant size were measured subsequently.

### Cloning and Plasmid Preparation

Full-length coding sequences of SmTRB (XP_002972715) were amplified from a total cDNA library generated from *Selaginella moellendorffii*. The PCR products were initially cloned into the pJET1.2/blunt vector using the CloneJET™ PCR Cloning Kit (Thermo Scientific; #K1231, #K1232) and verified by Sanger sequencing. The verified SmTRB inserts were recombined into pDONR221 and transferred into appropriate destination vectors using Gateway™ cloning technology (Invitrogen), including pGBKT7-GW, pGADT7-GW, pMDC7:i35S-GFP, and pMDC7:i35S-mCherry. The constructs pMDC7:i35S-PWO1-GFP, pMDC7:i35S-PWO1-mCherry, pMDC7:i35S-SmpWOa-GFP, pMDC7:i35S-SmpWOb-GFP, and dCas9:2×MS2:3×GFP were obtained from previously published sources (Khan et al., 2026; Hohenstatt et al., 2018; Khosravi et al., 2020). The coding sequences of TRB1, TRB2, and TRB3 (Kusová et al., 2023) were assembled into the pMDC85:2X35S-RFP vector using In-Fusion® Snap Assembly (Takara Bio) following the manufacturer’s instructions. All the cloning primers are listed in Table S5.

### Yeast Two-Hybrid Assays

Yeast two-hybrid (Y2H) interactions were assessed using the *Saccharomyces cerevisiae* strain AH109. For each assay, yeast cells were co-transformed with combinations of pGADT7-Gal4 activation domain (AD) and pGBKT7-Gal4 DNA-binding domain (BD) constructs, following the manufacturer’s instructions in the Yeast Protocols Handbook (Clontech, version 325, PR973283 21). After transformation, cells were plated on synthetically defined (SD) medium lacking leucine (Leu) and tryptophan (Trp) to select for the presence of both plasmids. For initial growth, SD medium was supplemented with adenine (Ade) and histidine (His) (SD/-Trp/-Leu/+Ade/+His) and incubated at 30°C for 2–3 days. Protein-protein interactions were evaluated on high-stringency selective media (SD/-Trp/-Leu/-Ade/-His) to assess the strength of interactions. Each assay included appropriate controls, consisting of co-transformations with empty AD and BD vectors, to rule out autoactivation or non-specific interactions. Assays were repeated with at least three independent colonies to confirm reproducibility of the interaction patterns.

### Subcellular Localization and FRET Assays

The pMDC7 (estradiol-inducible) or pMDC85 constructs tagged with GFP or mCherry were transformed into *Agrobacterium tumefaciens* strain GV3101 using the freeze–thaw method (Weigel and Glazebrook 2006). Following transformation, colonies were grown for 2 days on YEB plates (Sigma-Aldrich, Y1625) and cultured overnight in liquid YEB in 10 ml of LB. Cultures were sub-cultured in 50 mL LB for 3-4 hours at 28 °C with shaking at 180 rpm, then harvested and resuspended in infiltration medium containing 10 mM MgCl₂ (Sigma-Aldrich, M8266), 10 mM MES (pH 5.6; Sigma-Aldrich, M3671), and 200 μM acetosyringone (Sigma-Aldrich, D134406) to an OD₆₀₀ of 0.6, and incubated at room temperature for 1 hour. The bacterial suspensions were infiltrated into the abaxial side of 3-4-week-old *Nicotiana benthamiana* leaves using 2-mL syringes and incubated for 48-72 hours. Protein expression was induced by spraying the leaves with a solution of beta-estradiol (Sigma-Aldrich, E2758) in 0.1% (v/v) TWEEN-20 (Sigma-Aldrich, P1379). Leaf samples were analyzed 6-8 hours post-induction using a confocal microscope (Olympus FV3000; ZEISS LSM Airyscan), and images were processed using FIJI/ImageJ (Schindelin et al. 2012). For FLIM-FRET experiments, constructs were transiently expressed in *Nicotiana tabacum* (SR1 Petit Havana) leaf epidermal cells following the *Agrobacterium* infiltration method described by *Voinnet et al. (2000)* (Voinnet et al. 2000). Imaging was performed on a Zeiss LSM 780 AxioObserver equipped with an In Tune laser (488–640 nm, <3 nm width, pulsed at 40 MHz, 1.5 mW) and a C-Apochromat 63× water objective (NA 1.2). FLIM-FRET data were acquired using an HPM-100-40 hybrid detector (Becker & Hickl GmbH) with a Simple-Tau 150N TCSPC system and DCC-100 detector controller for photon counting. Image processing was performed with Zen 2.3 light edition (Zeiss), and FLIM data were analyzed using SPCM 64 v9.8 and SPCImage v7.3 (Becker & Hickl GmbH) employing a multiexponential decay model.

### Protein Structure Prediction and Visualization

Protein structures were predicted using the AlphaFold3 Protein Structure Database via https://alphafold.ebi.ac.uk (Abramson et al. 2024). For each protein, the server generated predicted three-dimensional 5 models along with confidence metrics, including per-residue pLDDT (predicted Local Distance Difference Test) scores and Predicted Aligned Error (PAE) plots. The PAE plots indicate the uncertainty in the relative positions of protein domains, with blue representing low uncertainty and red indicating higher uncertainty. The predicted structures were downloaded in PDB format and subsequently visualized using UCSF ChimeraX (Pettersen et al. 2021; Meng et al. 2023) for structural inspection.

### Chromatin Immunoprecipitation (ChIP) assay

ChIP assays were performed as previously described with minor modifications (Yang et al. 2024). Briefly, 10-day-old stably transformed *PWO1pro::PWO1-GFP/trb123* and *PWO1pro::PWO1-GFP/WT* seedlings (∼2.0 g per biological replicate) grown under long-day conditions were collected. For *PWO1pro::PWO1-GFP/trb123,* the material of half of each replicate was derived from the homozygous *trb1 trb2 trb3* triple mutant background (*PWO1pro::PWO1-GFP/trb1-2 trb2-2 trb3-1*), whereas the remaining material consisted of a mixture of *trb* double homozygous and two-homozygous/one-heterozygous plants (*PWO1pro::PWO1-GFP/trb1-2 trb2-2 trb3-1^+/−^*). Samples were crosslinked with 1% formaldehyde in MC buffer (10 mM potassium phosphate, pH 7.0, 50 mM NaCl, and 0.1 M sucrose) and quenched with 0.1 M glycine for 5 min under vacuum. Fixed tissue was ground in liquid nitrogen and extracted in lysis buffer containing 50 mM HEPES (pH 7.5), 150 mM NaCl, 1 mM EDTA, 0.1% deoxycholate, 0.1% SDS, 1% Triton X-100, 1 mM PMSF, and protease inhibitor cocktail. Chromatin was fragmented to <500 bp by sonication using a Bioruptor® Pico (Diagenode). Immunoprecipitation was performed using anti-GFP antibody (Abcam, ab290), and DNA–protein complexes were captured with Magna ChIP™ Protein A+G magnetic beads (Millipore, 16–663). Following reversal of crosslinks, DNA was purified by phenol–chloroform extraction followed by ethanol precipitation. Two biologically independent DNA libraries were constructed using VAHTS Universal Pro DNA Library Prep Kit for MGI (Vazyme Biotech Co.,Ltd) and sequenced using T7 PE150 platform.

### ChIP-seq Data Processing, Peak Annotation, and Functional Analysis

Published ChIP-seq raw data for proteins (PWO1-FLAG (Zheng et al., 2023), FLAG-TRB1, FLAG-TRB2, FLAG-TRB3 (Wang et al. 2023) and histone modifications in *Col-0* seedlings (∼12–14 DAG), including H3K27me3 (Wang et al. 2023), H3K4me3 (Wang et al. 2023), and H3K27ac (Chen et al. 2017), were downloaded from Sequence Read Archive (SRA) (Table S6). Paired-end reads were quality-checked using FastQC v0.11.9 (Andrews 2010) and adapters/low-quality sequences were removed with Trimmomatic v0.39 (TruSeq3-PE adapters) (Bolger et al. 2014). Reads were aligned to the *Arabidopsis thaliana* TAIR10 genome using Bowtie2 v2.4.5 (Langmead and Salzberg 2012). SAM alignments were converted to BAM, sorted, indexed, and filtered with SAMtools v1.10 (Li et al. 2009) to retain uniquely mapped reads (MAPQ ≥1), excluding unmapped and secondary alignments. Peak calling was performed with MACS2 v2.2.5 (Zhang et al. 2008) using biological replicates and corresponding input controls. PWO1 and TRBs peaks were identified as narrow sites, while histone modifications (H3K27me3, H3K4me3, H3K27ac) were analyzed as broad enrichment domains. Genome-wide coverage tracks were generated with deepTools v3.3.1 (Ramírez et al. 2016) using Reads Per Kilobase of transcript per Million mapped reads (RPKM) normalization for visualization. BED coordinates of identified peaks were annotated against TAIR10 gene models using rtracklayer, GenomicRanges, and ChIPseeker (Lawrence et al. 2013; Yu et al. 2015), with promoters defined as −1000 bp to +100 bp relative to transcription start sites. Other features (exons, introns, UTRs, intergenic regions) were annotated according to gene models. Unbiased k-means clustering of signal matrices was applied to partition peaks into clusters, and cluster-specific gene lists were extracted. Functional enrichment analyses (e.g., Gene Ontology) were performed using clusterProfiler (Yu et al. 2012) and org.At.tair.db (Carlson 2019) (BH correction, q ≤ 0.05), and feature distributions and enrichment results were visualized using ggplot2 (Wickham 2016). The bigWig (.bw) files for replicate 1 of TRB1-del-Myb-FLAG and TRB1-del-GH1(H1/5)-FLAG datasets (Wang et al. 2025) were obtained from Gene Expression Omnibus (GEO) and directly analyzed using deepTools to generate metagene profiles. Published chromatin state annotations for Arabidopsis were obtained from the 26-state segmentation model (Shukla et al. 2026). ChIP-seq peak coordinates were intersected with chromatin state segments using GenomicRanges, and the proportion of peaks overlapping each chromatin state was calculated for comparative analysis and visualization.

### Histochemical Analysis

To visualize lignin deposition in the main stem, stems from 28-54-day-old Arabidopsis plants were collected, and transverse sections from the basal region of the stem were manually prepared using a razor blade. Sections were incubated in 50% ethanol for 15 min and subsequently stained with 0.01% Basic Fuchsin (Sigma-Aldrich, 857343) for 15 min at room temperature with gentle shaking, as previously described with slight modifications (Kapp et al. 2015). After staining, sections were washed three times with distilled water (5 min each) to remove excess dye and then imaged using a fluorescence microscope with 4× and 10× objectives.

### Gene Expression Analyses

Total RNA was extracted from 14-day-old seedlings and 28-day-old stems after bolting but before flowering using the Thermo Scientific RNA extraction kit, following the manufacturer’s instructions. First-strand cDNA was synthesized from 1 µg of total RNA using the RevertAid First Strand cDNA Synthesis Kit (Thermo Scientific). The resulting 20 µL cDNA was diluted 1:20 and vortexed prior to use as a template. Reactions were performed using SYBR Green fluorescent dye (ROX SYBR® MasterMix blue dTTP, Takyon™) in 384-well plates, with each sample analyzed in triplicate. Master mixes were prepared individually for each target gene, and 4 µL of diluted cDNA was combined with 6 µL of qPCR master mix per well. Plates were sealed and loaded into a QuantStudio™ 5 Real-Time PCR System (Thermo Fisher Scientific). TIP41 was used as a housekeeping control for normalization.

## Data Statement

Sequence information for the genes analyzed in this study is available in the NCBI, TAIR10, and UniProt databases under the following accession numbers: *At3g03140 (PWO1), At1g49950 (TRB1), At5g67580 (TRB2), At3g49850 (TRB3), and XP_002972715 (SmTRB)*.

## Acknowledgments

We sincerely thank Prof. Dr. Andreas Houben and Dr. Solmaz Khosravi (Leibniz Institute of Plant Genetics and Crop Plant Research (IPK), Germany) for providing the dCas9:2×MS2:GFP construct for telomere visualization in *N. benthamiana*. We are grateful to Cordula Braatz and Ruth Lintermann for technical assistance, and Marina Hasse for organizational and administrative support. We thank Dr. Franziska Turck (Max Planck Institute for Plant Breeding Research, Germany) for providing *trb’s* double mutant seeds, and Prof. Steven E. Jacobsen (University of California, USA) for providing *trb1 trb2 trb3* segregating seeds. We thank Mohan Govindasamy, University of Galway for valuable discussions and assistance with ChIP-seq data analysis. We would like to acknowledge the assistance of the Core Facility BioSupraMol supported by the DFG and the infrastructure provided by the Research Building SupraFAB as well as the DFG for the funding of the Zeiss LSM 980 with AiryScan2 and PicoQuant FLIM (DFG project number 503201387)

## Author contributions

AK, BG, IM, SF and DS conceptualized the experimental approach and designed the methodology. AK performed the majority of molecular biology experiments, including construct generation, protein localization, Arabidopsis mutant generation via crossing, confocal microscopy analyses, ChIP-seq analysis, AF3-modeling, and data visualization, and manuscript writing. AKu and JS carried out the FLIM-FRET experiments. BG analyzed the preliminary genomic co-occupancy/shared chromatin association between PWO1 and TRB1, protein co-localization experiments and confocal microscopy analyses (as part of his doctoral thesis), and discussed the results. ALVK & LH performed genotypic and phenotypic experiments for *pwo1-trb’s* mutans. All authors contributed to the interpretation of the results. AK, DS, KCSP, IM, SF, PPS, and JH revised the manuscript.

