## Supplementary Figures for "PWO1 and TRB proteins coordinate chromatin regulation to prevent premature differentiation and ectopic lignin deposition in Arabidopsis"

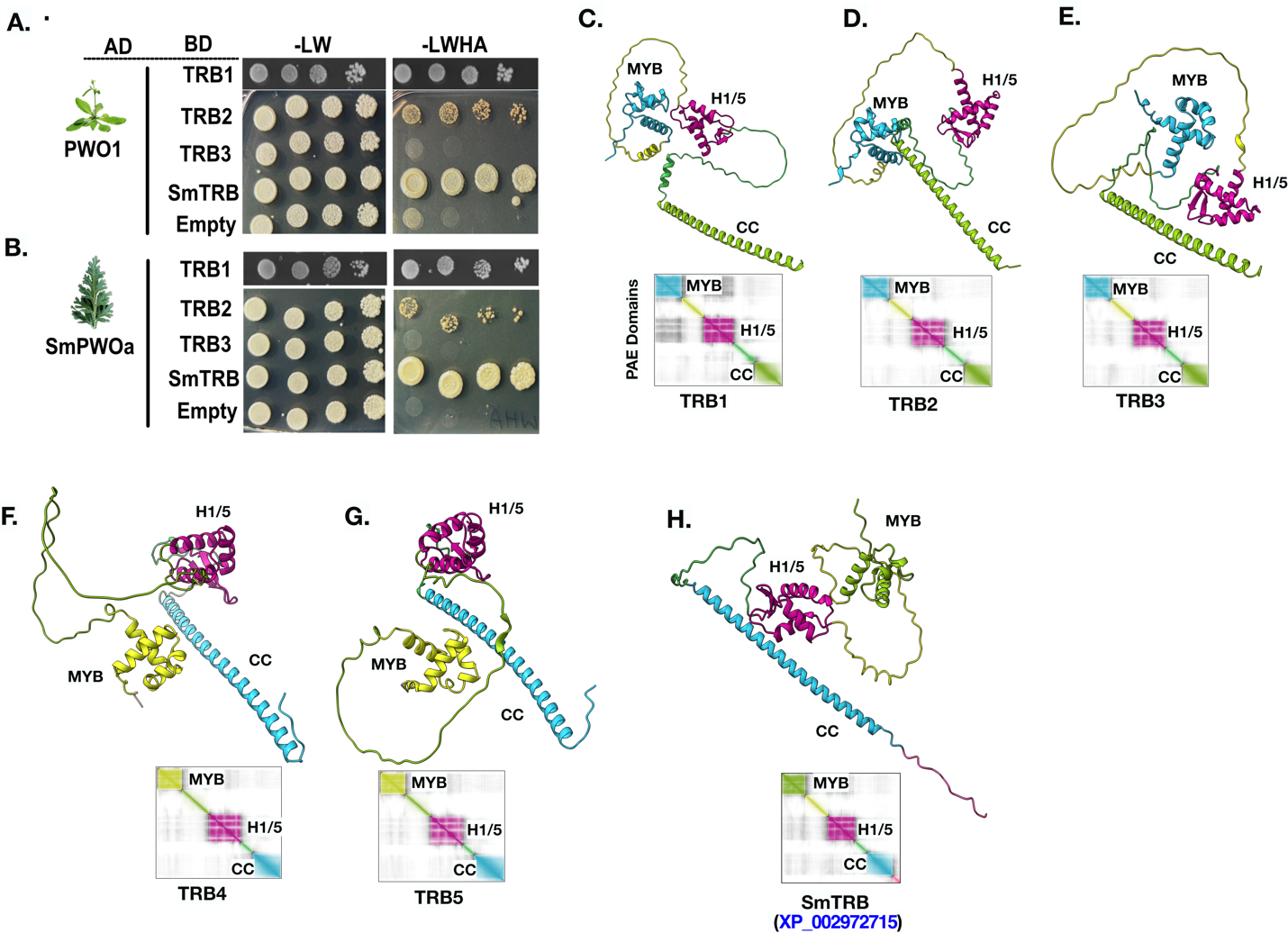

**Figure S1. PWO-TRB interactions and structural conservation of TRB proteins across evolution.** Yeast two-hybrid (Y2H) analyses of **A.** PWO1 (Arabidopsis) and **B.** SmPWOa (*S. moellendorffii*) interaction with TRB1-3 and SmTRB. The interactions were evaluated on non-selective medium (−LW; lacking leucine and tryptophan), to facilitate plasmid co-transformation, and on selective medium (−LWAH; lacking leucine, tryptophan, adenine, and histidine), to test the interactions. AF3-based structure predictions and associated PAE domain plots for **C-G.** *Arabidopsis* TRB1-5, and **H.** *S. moellendorffii* SmTRB (XP_002972715). The color of each domain within 3D-structure of TRB proteins corresponds to the colors in PAE plot.

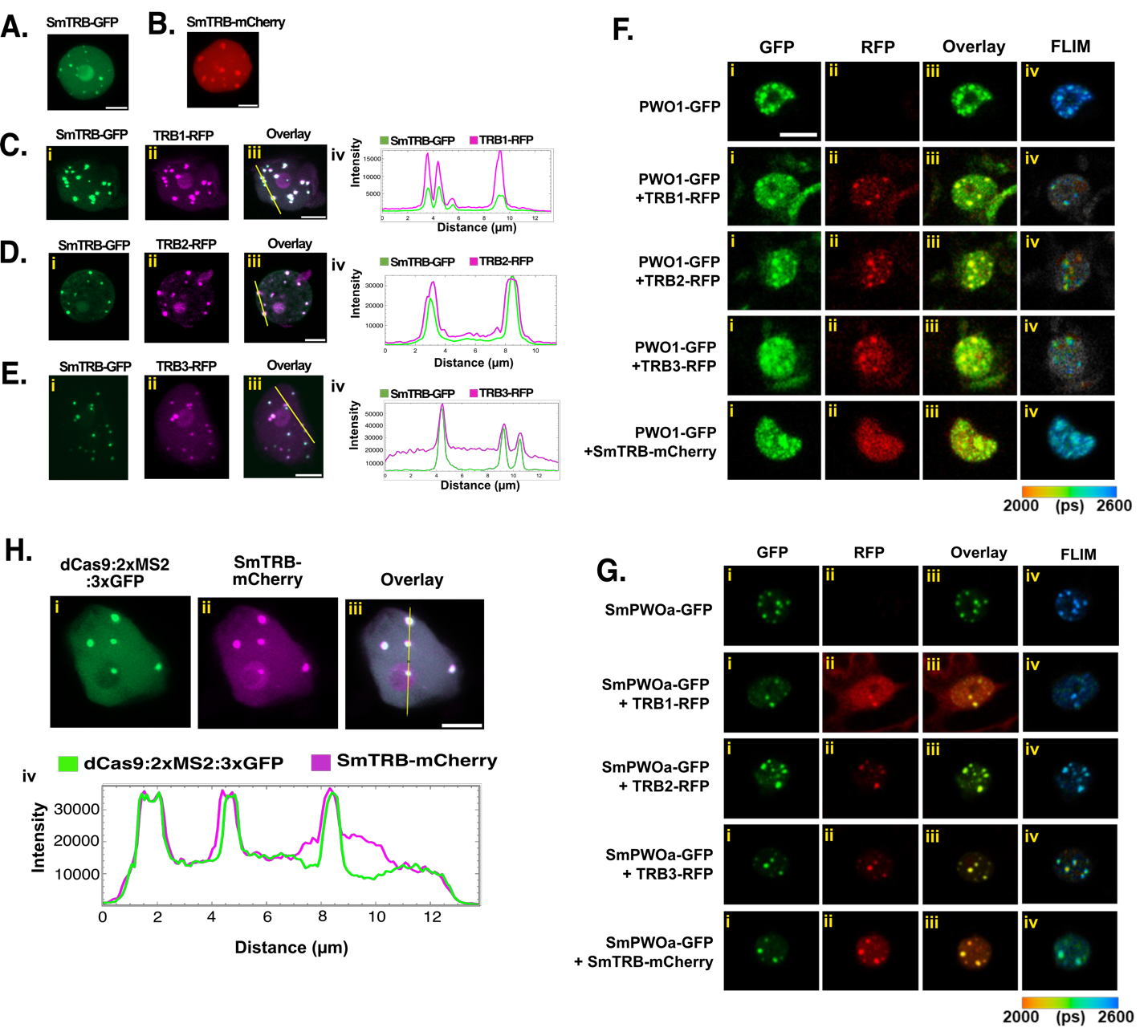

**Figure S2: SmTRBs co-localization with Arabidopsis TRB1-3 and FLIM-FRET confocal imaging of PWO1/SmPWOa and TRBs/SmTRB interactions.** Representative confocal microscopy images of *N. benthamiana* leaf nuclei infiltrated with **A.** *i35S::SmTRB-GFP* and **B.** *i35S::SmTRB-mCherry*. **C-E.** *i35S::SmTRB-GFP* was co-infiltrated with **C_i–iii:** *35S::TRB1-RFP*, **D_i–iii:** *35S::TRB2-RFP*, and **E_i–iii:** *35S::TRB3-RFP*. For each co-infiltration, fluorescence intensity profiles of GFP and RFP along the yellow line are shown in **C-E_iv**. Scale bar = 5 µm. (*SmTRB; XP_002972715*). Fluorescence lifetime images (FLIM) were acquired using a FLIM-FRET assay for *i35S::PWO1-GFP* **F.** and *i35S::SmPWOa-GFP* **G.** co-expressed with *35S::TRB1-RFP, 35S::TRB2-RFP, 35S::TRB3-RFP*, and *i35S::SmTRB-mCherry* in transiently transformed *N. tabacum* leaves. FLIM–FRET data are presented as false-color images using a fluorescence lifetime look-up table (LUT), where color represents fluorescence lifetime. Images represent results quantified in Figure 1H-I. Scale bar = 10 μm.

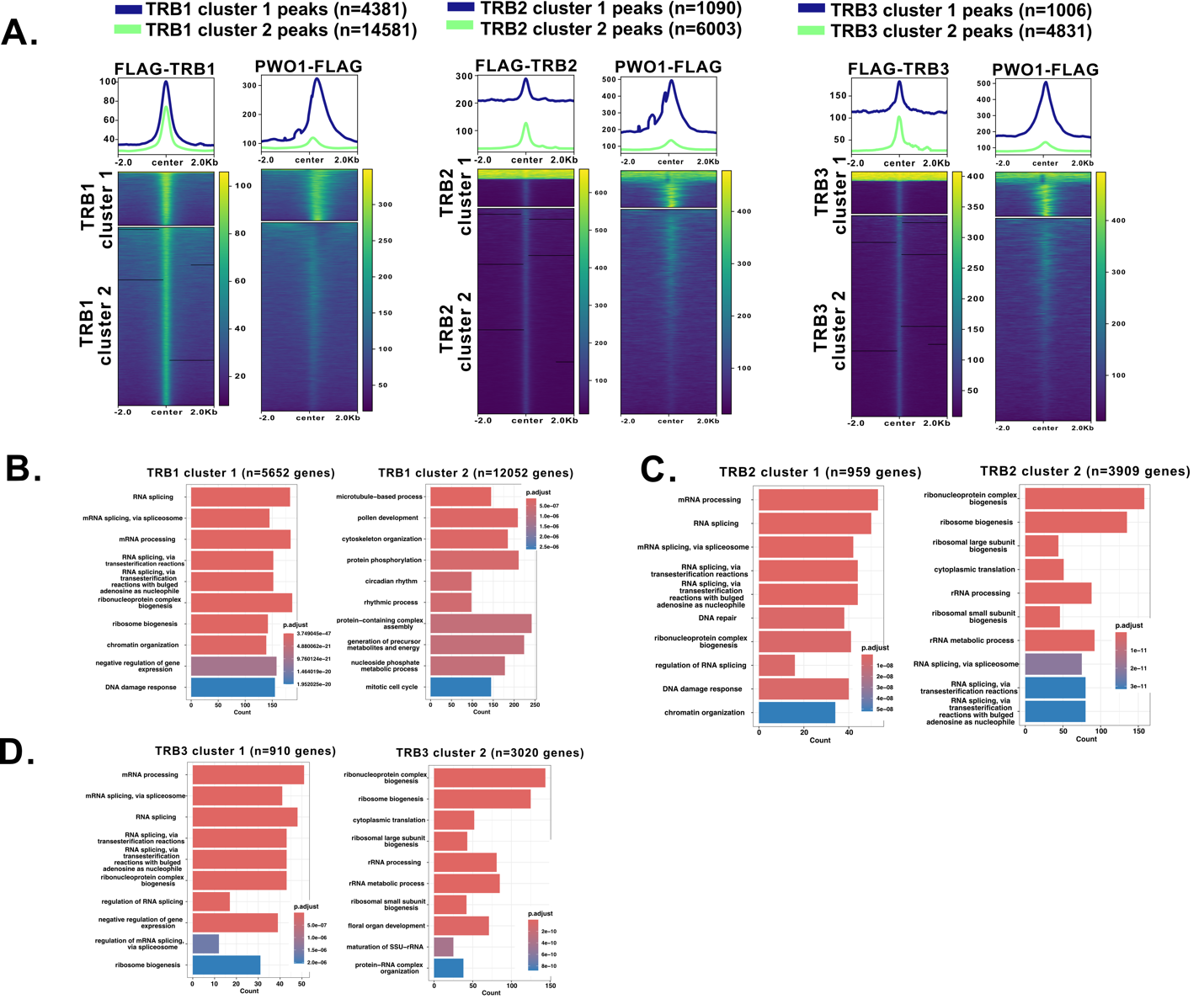

**Figure S3. PWO1 peak enrichment over TRB targets and functional annotation.**
**A.** Metaplots and heatmaps showing normalized ChIP-seq signals of PWO1-FLAG (Zheng *et al.*, 2023) enrichment over FLAG-TRB1, FLAG-TRB2, and FLAG-TRB3 peaks (Wang *et al.*, 2023). TRB peaks were classified into two clusters based on PWO1 enrichment. **B-D**. Gene Ontology (GO) enrichment analysis (Biological Process, BP) of genes associated with TRB1-3 clusters 1 and 2. Bars represent the top 10 significantly enriched terms, with lengths proportional to the number of associated genes. Significance was determined using clusterProfiler (Benjamini–Hochberg adjusted P < 0.05).

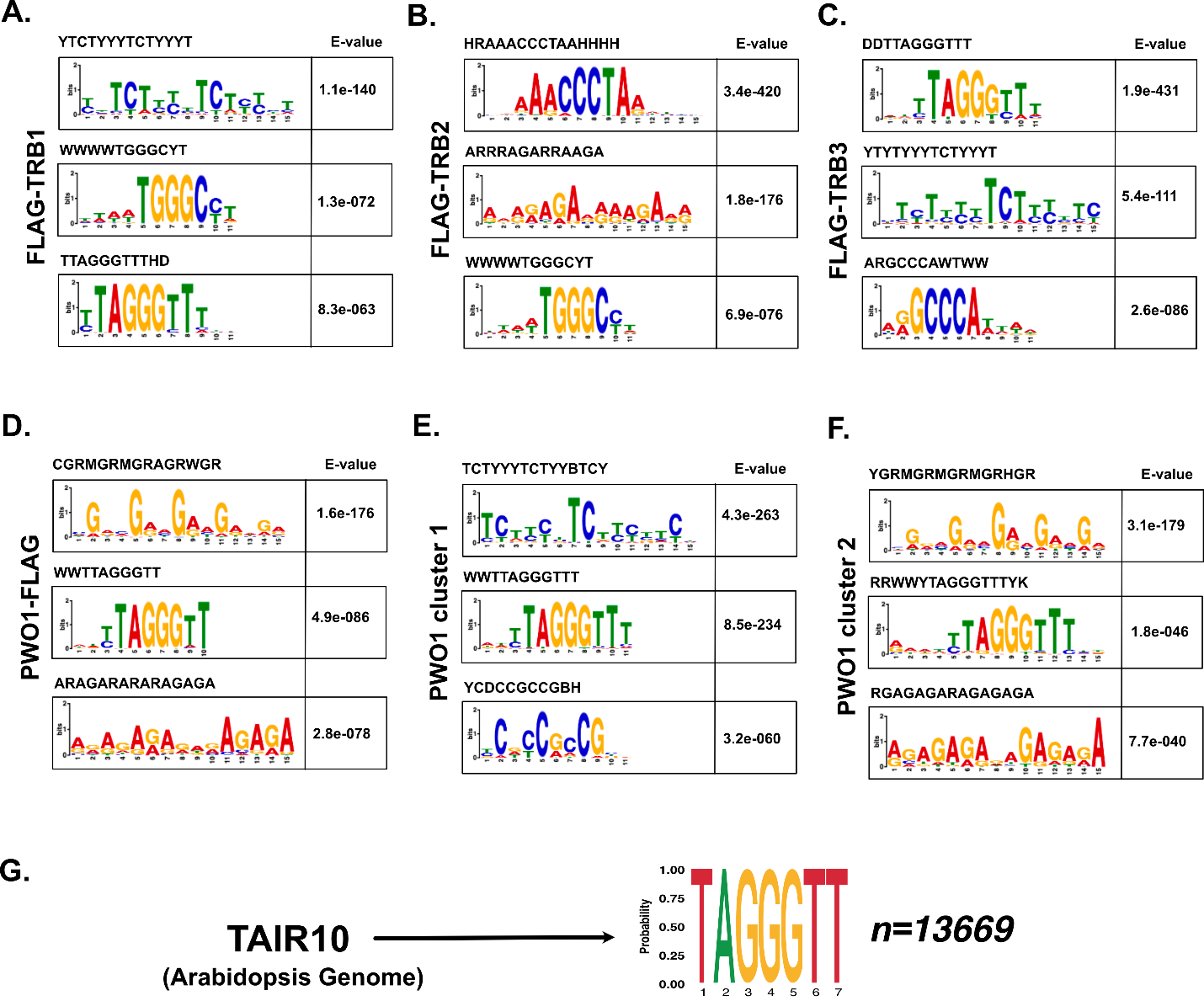

**Figure S4: DNA sequence motifs enriched at PWO1 and TRB binding sites.**
Three topmost enriched DNA sequence motifs identified for **A.** FLAG-TRB1, **B.** FLAG-TRB2, and **C.** FLAG-TRB3, **D.** PWO1-FLAG, **E.** PWO1 cluster 1 and **F.** PWO1 cluster 2 using MEME-ChIP analysis (https://meme-suite.org/meme/tools/meme-chip). Motif logos represent the consensus sequences, with the height of each letter proportional to nucleotide conservation. E-values indicate statistical significance, with lower values reflecting higher enrichment. **G.** Genomic occurrences of the TAGGGTT *telo*-box motif in the TAIR10 genome. **A-F.** For each motif enrichment, the consensus sequence is shown on the top of the logo.

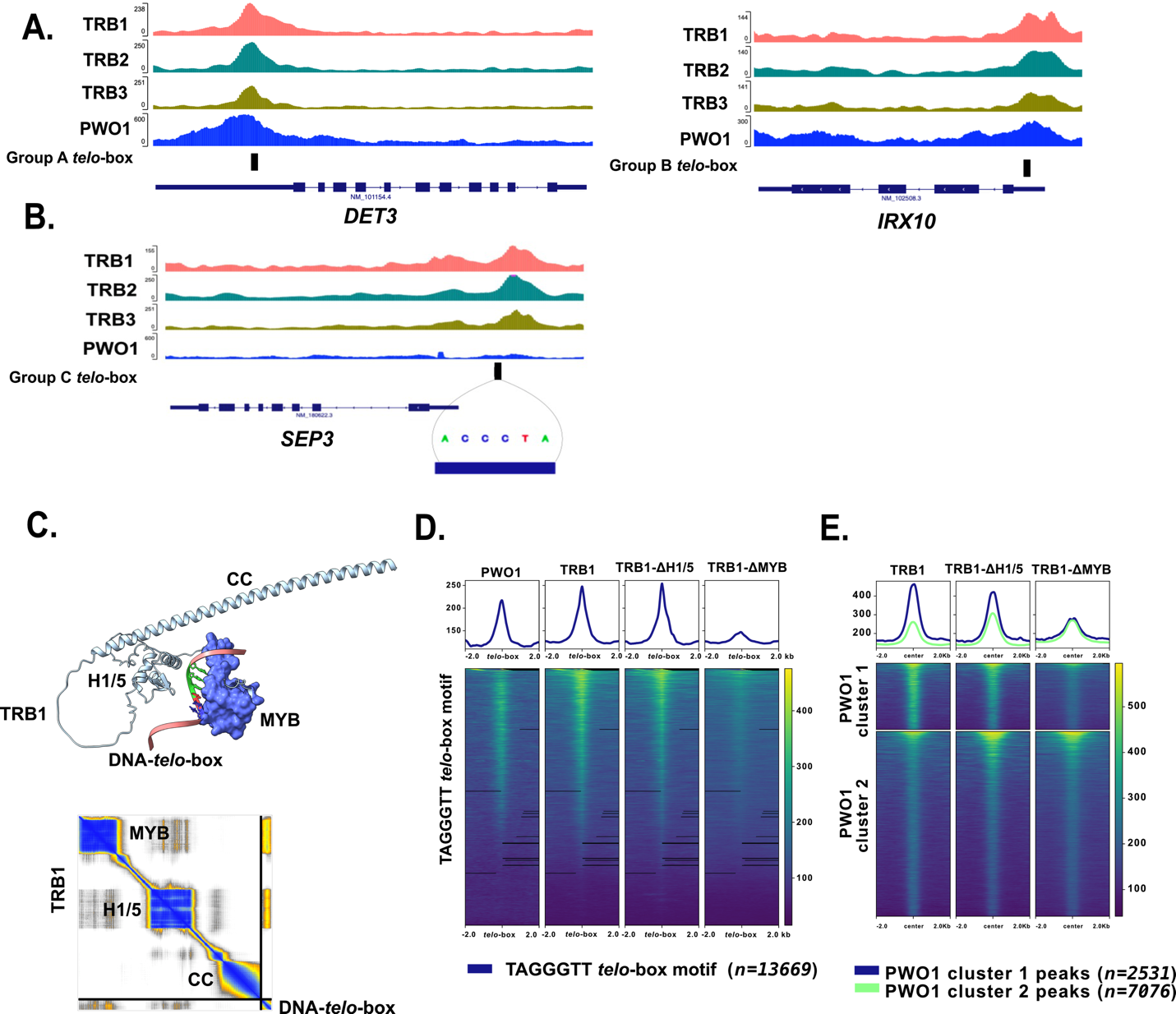

**Figure S5: Binding of TRB1 variants with deleted MYB or H1/5 domains over *telo-box* motifs as well as co-shared PWO1-TRBs targets. A-B.** Integrative Genomics Viewer (IGV) screenshots showing ChIP-seq signals for PWO1-FLAG (Zheng *et al.*, 2023) and FLAG-TRBs (Wang *et al.*, 2023) binding at *telo*-box Group 1-3 over the representative genes such as **A.** *DE-ETIOLATED 3 (DET3), IRREGULAR XYLEM 10* *(IRX10), and* **B.** *SEPALLATA 3* (*SEP3*) genes. **C.** AlphaFold3 prediction of TRB1 binding to the *telo*-box motif in 3D space, with the PAE plot showing the confidence of the predicted domain-DNA interactions. **D-E.** Metaplots and heatmaps showing normalized ChIP-seq signals of PWO1 and replicate 1 of TRB1, TRB1-ΔH1/5, and TRB1-ΔMYB over (Wang *et al.*, 2025) **D.** TAGGGTT t*elo*-box motifs and **E.** PWO1 cluster 1 and cluster 2 regions.

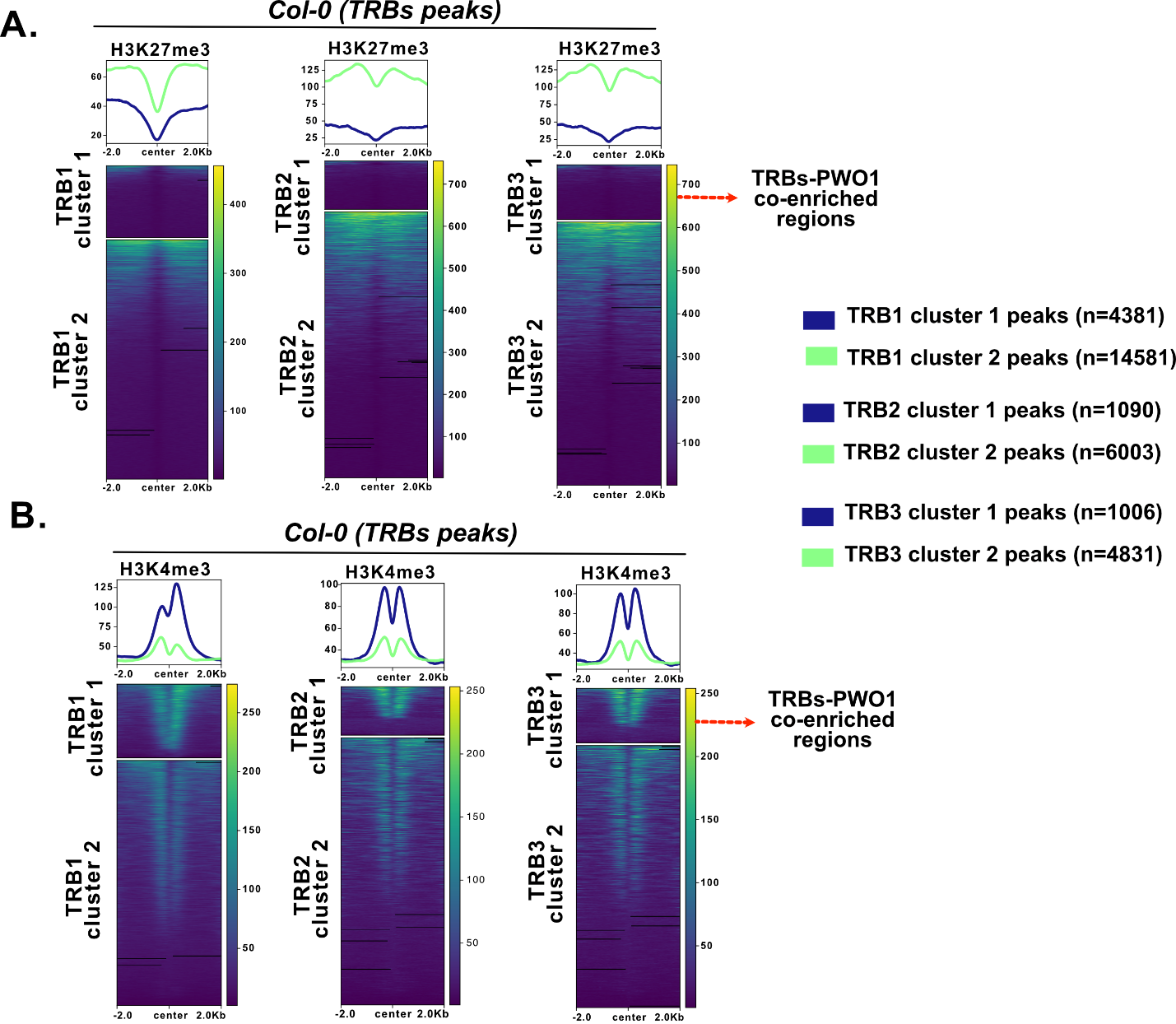

**Figure S6. H3K27me3 and H3K4me3 enrichment over TRB peaks. A-B.** Metaplots and heatmaps showing normalized ChIP-seq signals (Wang *et al.*, 2023) of **A.** H3K27me3 and **B.** H3K4me3 across TRB1-3 clusters 1 and 2, defined based on PWO1 enrichment (Figure S3). The red arrow indicates cluster 1, which is enriched for PWO1-TRB co-binding.

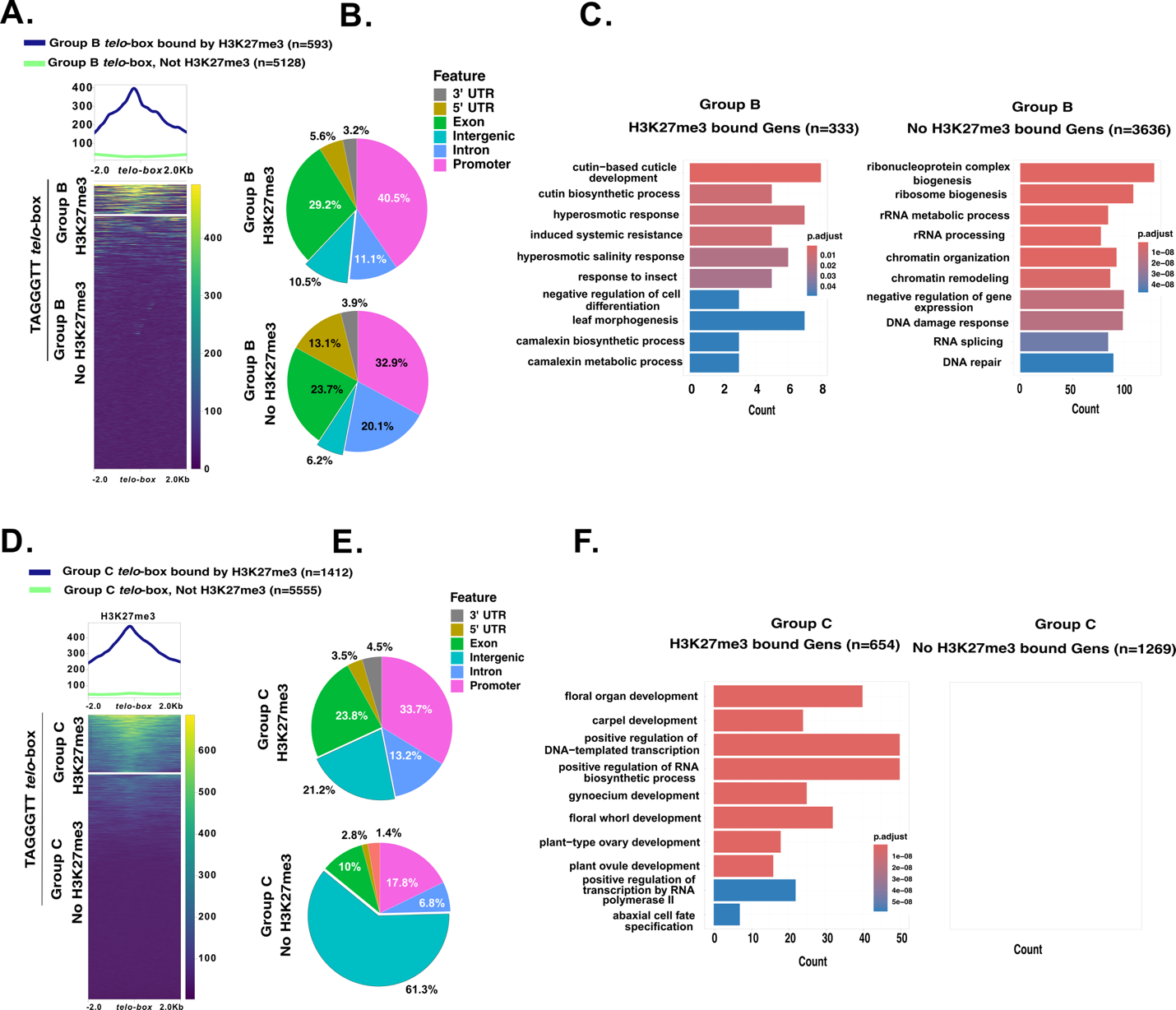

**Figure S7. H3K27me3 enrichment over Group A and B *telo*-box motif. A.** Metaplots and heatmaps showing normalized ChIP-seq signals (Wang *et al.*, 2023) of H3K27me3 over Group B *telo*-box motifs. **B.** Pie charts depicting the genomic annotation of Group B *telo*-box regions bound and unbound by H3K27me3, with **C.** GO enrichment analysis. **D.** Metaplots and heatmaps showing normalized ChIP-seq signals (Wang *et al.*, 2023) of H3K27me3 over Group C *telo*-box. **E.** Pie charts depicting the genomic annotation of Group C *telo*-box regions bound and unbound by H3K27me3, with **F.** GO enrichment analysis. Bars represent the top 10 significantly enriched biological processes, with lengths proportional to the number of genes associated with each term. Significance was determined using clusterProfiler (Benjamini–Hochberg adjusted p < 0.05).

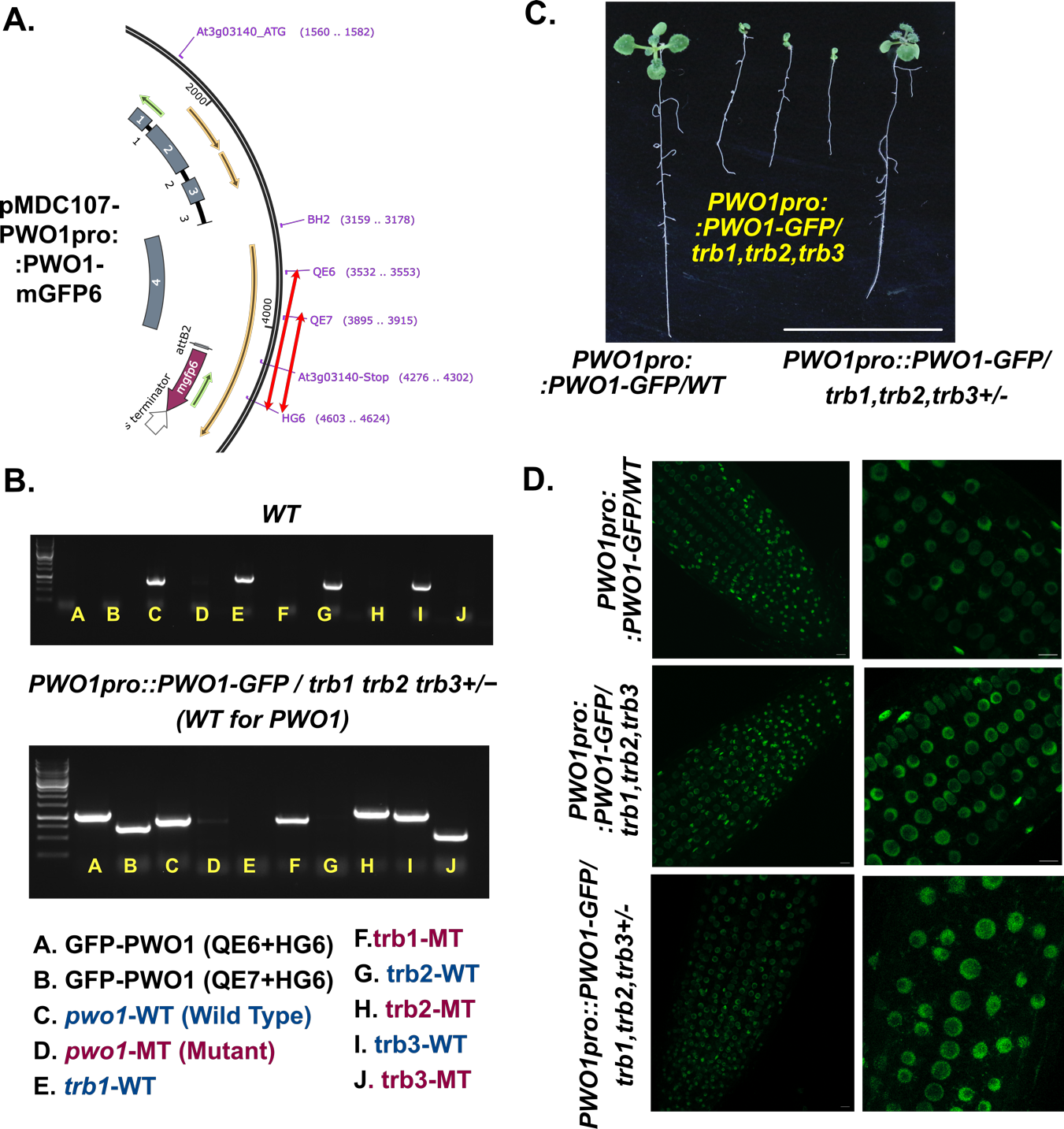

**Figure S8. Generation of *PWO1pro::PWO1-GFP* lines in the *trb1-2 trb2-2 trb3-1* background. A.** Schematic representation of the plasmid construct *pMDC107-PWO1pro::PWO1-mGFP6* expressing line used for crossing with the *trb1-2^+/−^ trb2-2 trb3* background (Wang *et al.*, 2023). Red arrows indicate the primers used to genotype the C-terminal PWO1-GFP fusion. **B.** Genotyping results and primer pairs used for PCR amplification in WT and *PWO1pro::PWO1-GFP / trb1-2 trb2-2 trb3* lines in a PWO1 WT background. **C.** Representative images of 2-week-old seedlings of *PWO1pro::PWO1-mGFP6*, *PWO1pro::PWO1-mGFP6* / *trb1-2 trb2-2 trb3*, and *PWO1pro::PWO1-mGFP6 / trb1-2 trb2-2 trb3+/−* in a PWO1 WT background. **D.** Confocal microscopy images of root tips of *PWO1pro::PWO1-GFP* expressed in WT, *trb1-2 trb2-2 trb3*, and *trb1-2 trb2-2 trb3+/−* backgrounds. Scale bar: 100 µm.

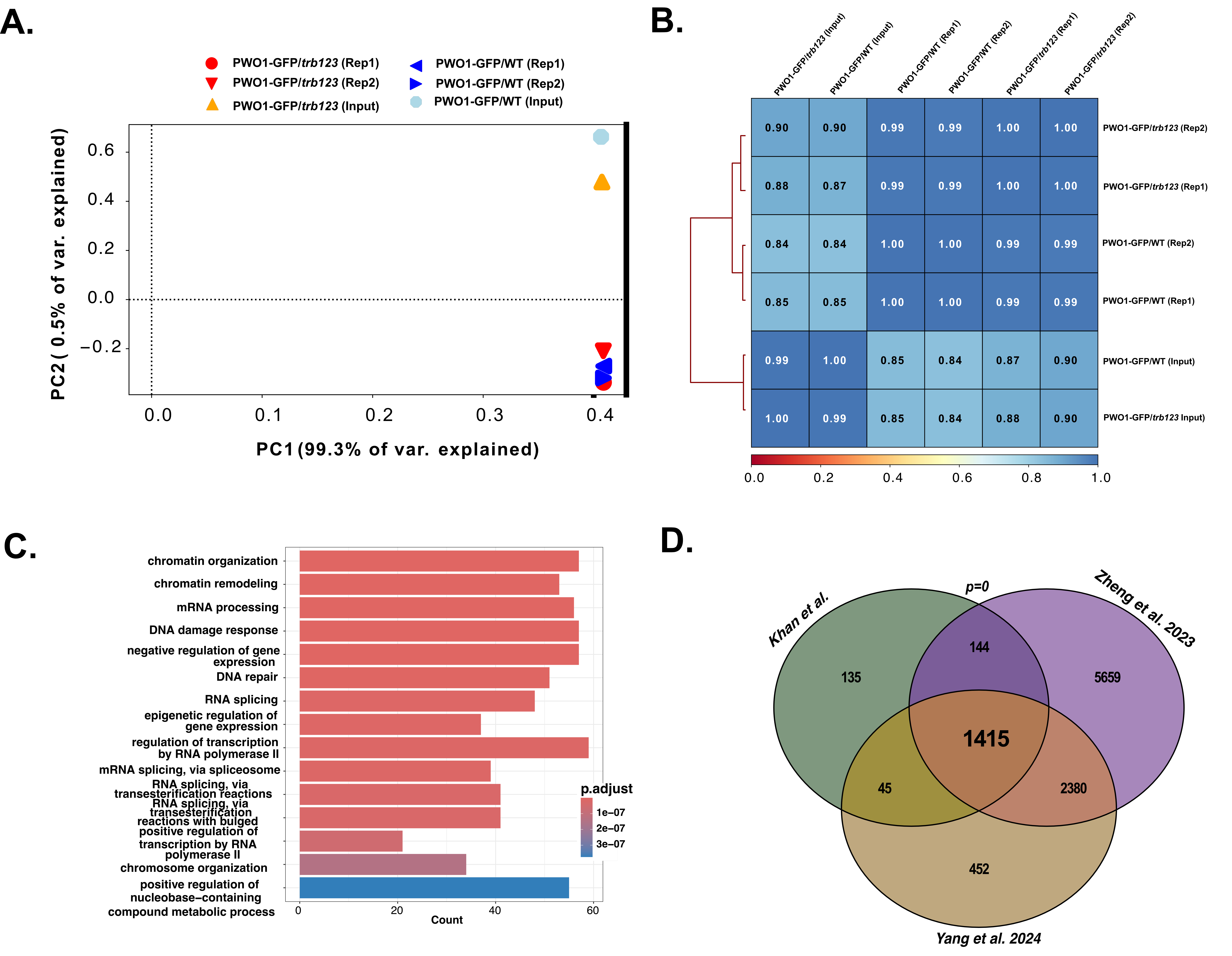

**Figure S9.** **Quality control and comparative analysis of ChIP-seq replicates.** **A.** Principal Component Analysis (PCA) showing the similarity and variation among biological replicates of PWO1pro::PWO1-GFP in the WT and *trb1 trb2 trb3* mutant backgrounds (a pool of *PWO1pro::PWO1-GFP/trb1-2 trb2-2 trb3-*1 and *PWO1pro::PWO1-GFP/trb1-2 trb2-2 trb3-1^+/−^;* referred to as the *trb123* mutant background). **B.** Pairwise correlation matrix based on Pearson correlation coefficients for all combinations of PWO1-GFP samples in WT and *trb123* mutant backgrounds. Analyses in A-B were performed using uniquely mapped BAM reads. **C.** Gene Ontology (GO) enrichment analysis of genes BP associated with PWO1-GFP in WT (*n = 1551*). Bars represent the top 10 significantly enriched terms, with bar length proportional to the number of associated genes. Significance was assessed using clusterProfiler with Benjamini-Hochberg adjusted P < 0.05. **D.** Venn diagram illustrating the overlap of PWO1-bound genes in WT across this study and previously published datasets (Zheng *et al.*, 2023; Yang *et al.*, 2024), showing a high degree of concordance.

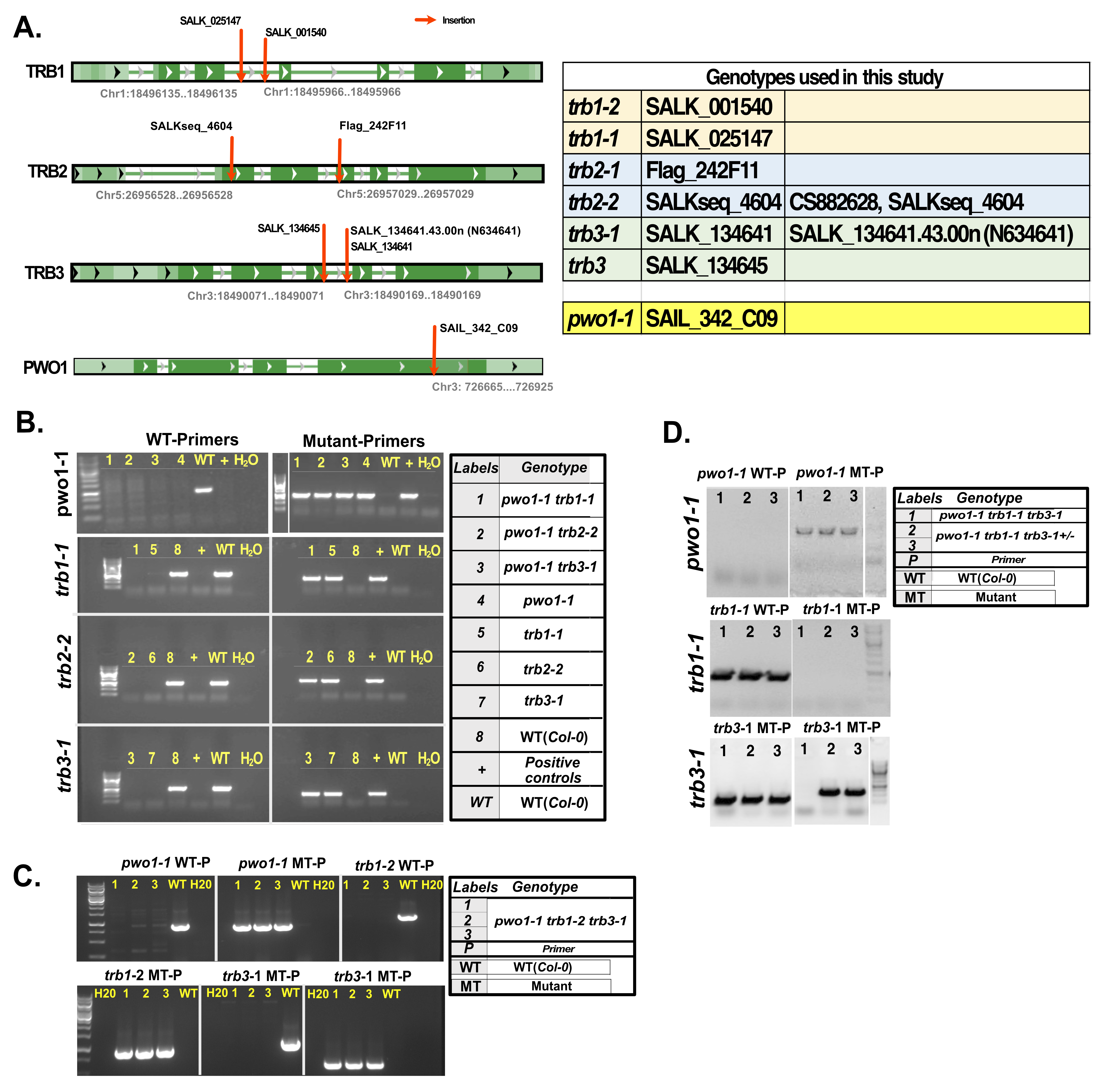

**Figure S10.** **Allelic combinations and genotyping PCR for *pwo1* and *trb* mutants.
A.** Genomic locations of all *trb* and *pwo1* mutant alleles shown via genome browser tracks, with chromosomal positions displayed for each mutant allele using the NCBI Genome Browser ([https://www.ncbi.nlm.nih.gov](https://www.ncbi.nlm.nih.gov/)). **B-D.** Genotyping PCR for **B.** *pwo1-trb’s* double mutants (*pwo1-1 trb1-1, pwo1-1 trb2-2, pwo1-1 trb3-1*), **C.** *pwo1-1 trb1-2 trb3-1*, and **D.** *pwo1-1 trb1-1 trb3-1* and *pwo1-1 trb1-1 trb3-1+/-* triple mutants.

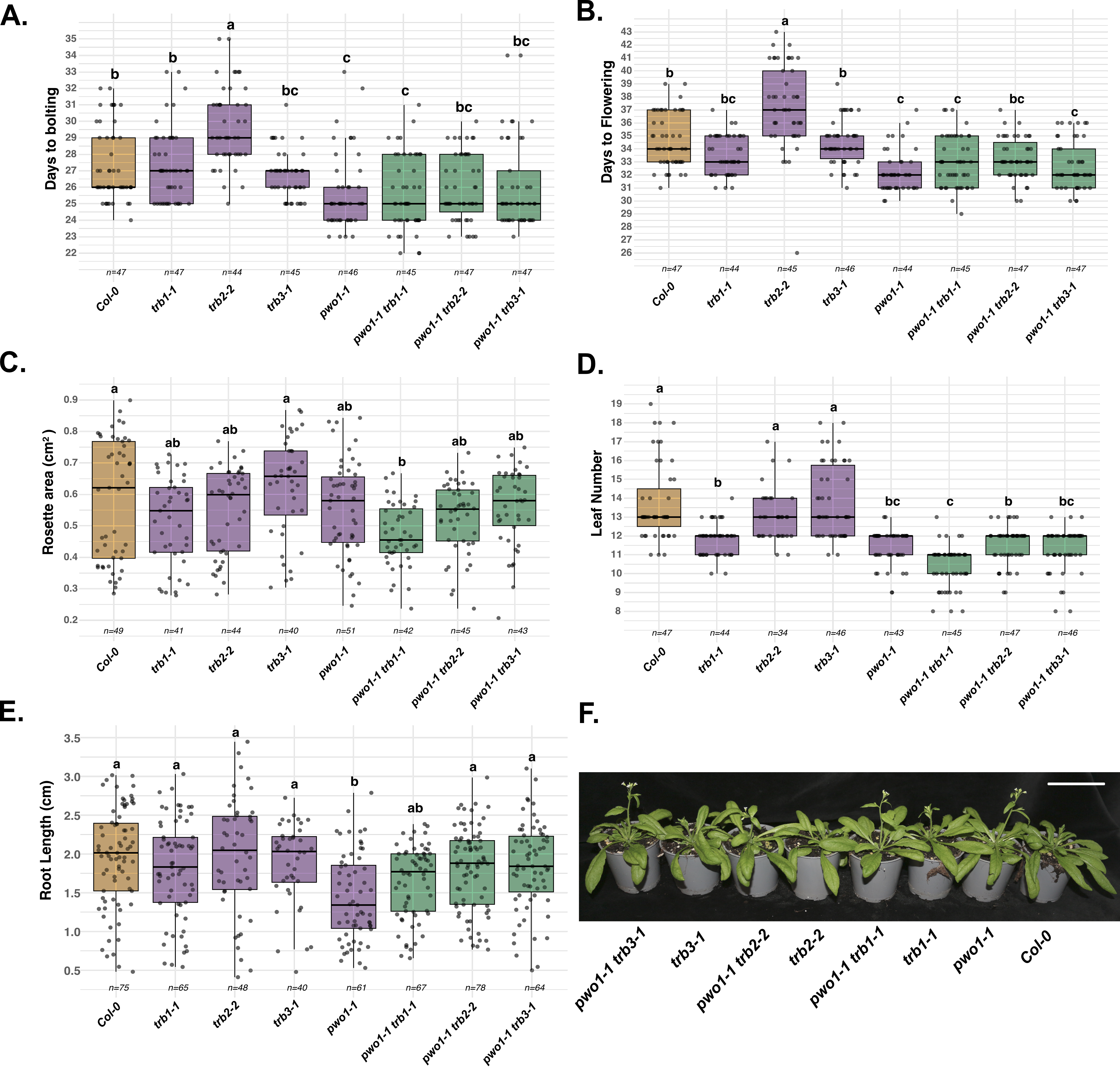

**Figure S11.** **Phenotypic analysis of *pwo1* and *trb* combinatorial double mutants in Arabidopsis plants.** Boxplots show the distribution of **A.** days to bolting, **B.** days to flowering, **C.** rosette area, **D.** leaf number at bolting, and **E.** root length for each genotype. Black points represent individual plants and are displayed with horizontal jitter. Box colors indicate genotype groups: orange for wild type (Col-0), purple for single mutants (*pwo1-1, trb1-1, trb2-2, trb3-1*) and green for double mutants (*pwo1-1 trb1-1, pwo1-1 trb2-2, pwo1-1 trb3-1*). Different letters above the boxes represent statistically significant differences between genotypes (one-way ANOVA followed by Tukey’s HSD test, *p* < 0.01). **F.** Representative pictures of genotypes (*pwo1-1 trb3-1, trb3-1, pwo1-1 trb2-2, trb2-2, pwo1-1 trb1-1, trb1,* and Col-0) at 31 days after germination. Scale bar = 5 cm.

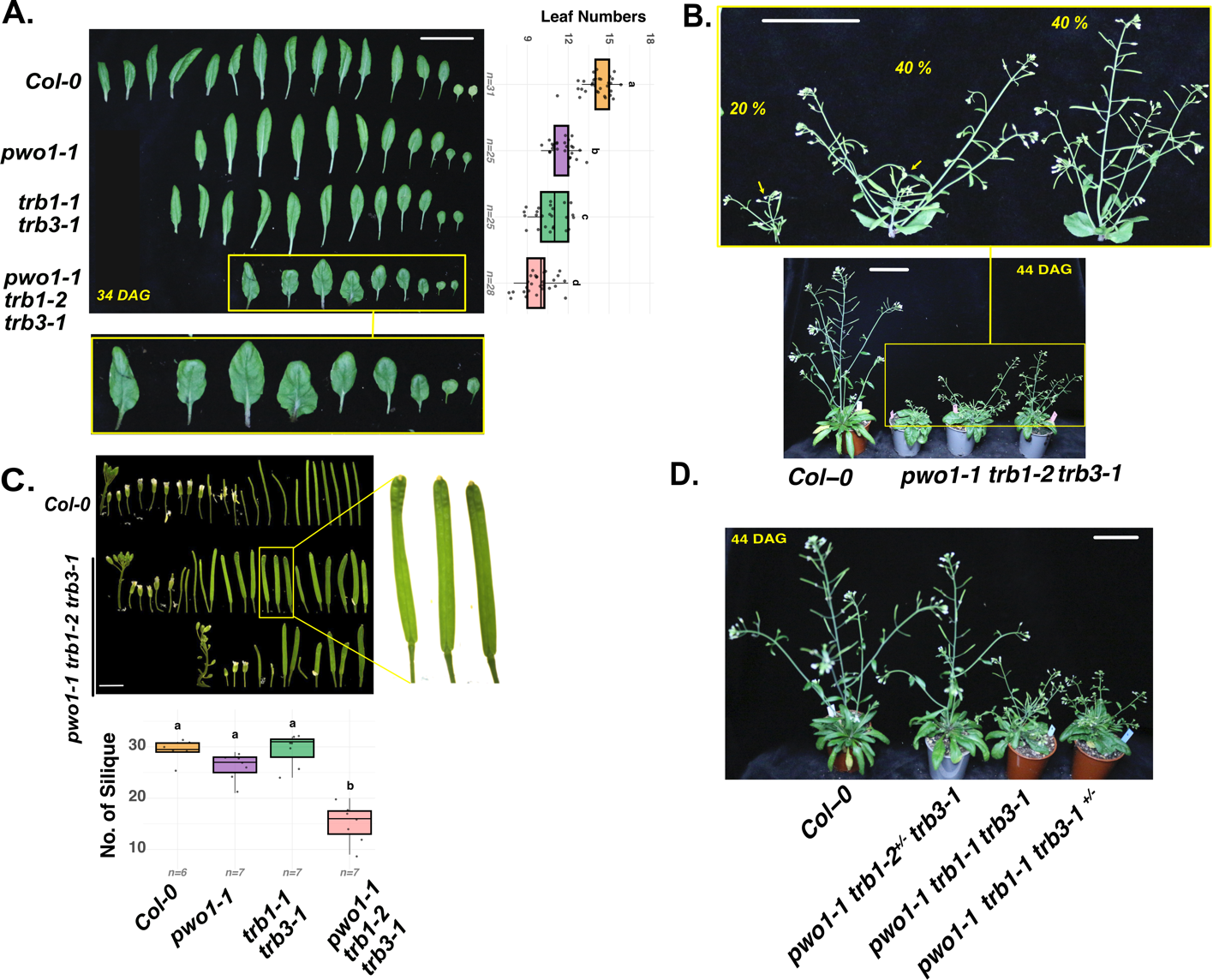

**Figure S12. Phenotypic analysis of *pwo1 trb1 trb3* triple mutant plants.** **A.** Leaf phenotypes of excised leaves from 34-day-old plants; **B.** main stem phenotypes observed in *pwo1-1 trb1-2 trb3-1* triple mutants, including complete main stem collapse shortly after bolting (20%); severely reduced main stem elongation accompanied by dominance of axillary branches (40%), and a short but morphologically intact main stem exhibiting irregular silique distribution (40%). Yellow arrow shows the main stem. **C.** Silique phenotypes of *pwo1-1 trb1-2 trb3-1* triple mutants compared with Col-0 and Number of siliques in main stem on 46 old plants. **D.** Phenotypes of *trb1-2* and *trb3-1* segregating lines in the *pwo1 trb1 trb3* mutant background with homozygous *pwo1-1 trb1-1 trb3-1* compared with Col-0 plants. In panel A, Box colors indicate genotype groups. Different letters above the boxes indicate statistically significant differences between genotypes (one-way ANOVA followed by Tukey’s HSD test, *p* < 0.01). Scale bar = 5 cm.

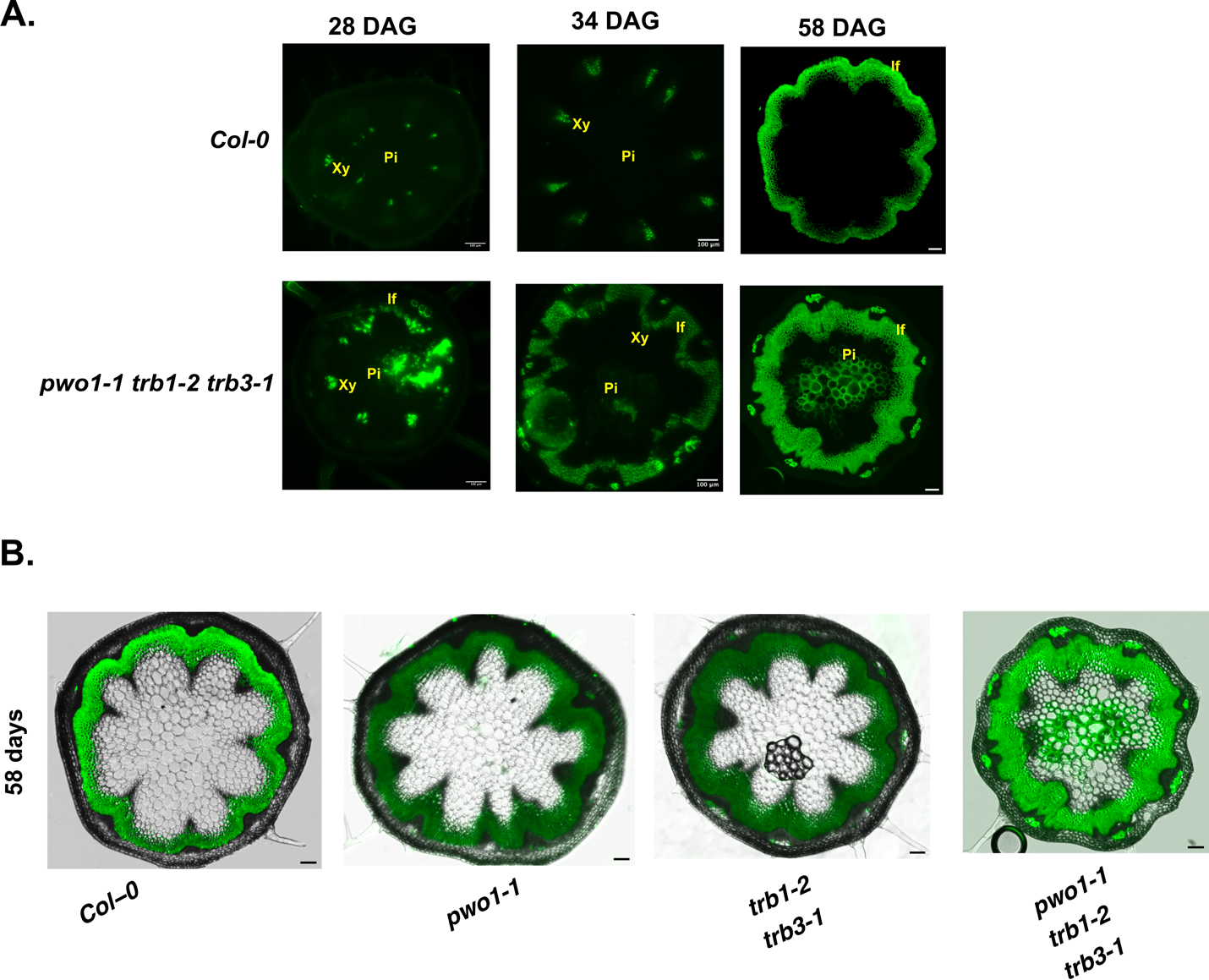

**Figure S13. Ectopic lignification in *pwo1 trb1 trb3* triple mutant. A.** Transverse sections (TS) taken from the base of the main stem of 58-day-old *pwo1, trb1 trb3, pwo1 trb1 trb3*, and Col-0 plants, stained with Basic Fuchsin. **B.** TS of the basal stem at 28 days, showing only Basic Fuchsin–stained lignified regions. **C.** Transverse sections at 34 days for Col-0, *pwo1 trb1-2 trb3-1*, and *pwo1-1 trb1-2^+/-^ trb3-1*, highlighting Basic Fuchsin-stained interfascicular fibers. X = xylem; Pi = pith.

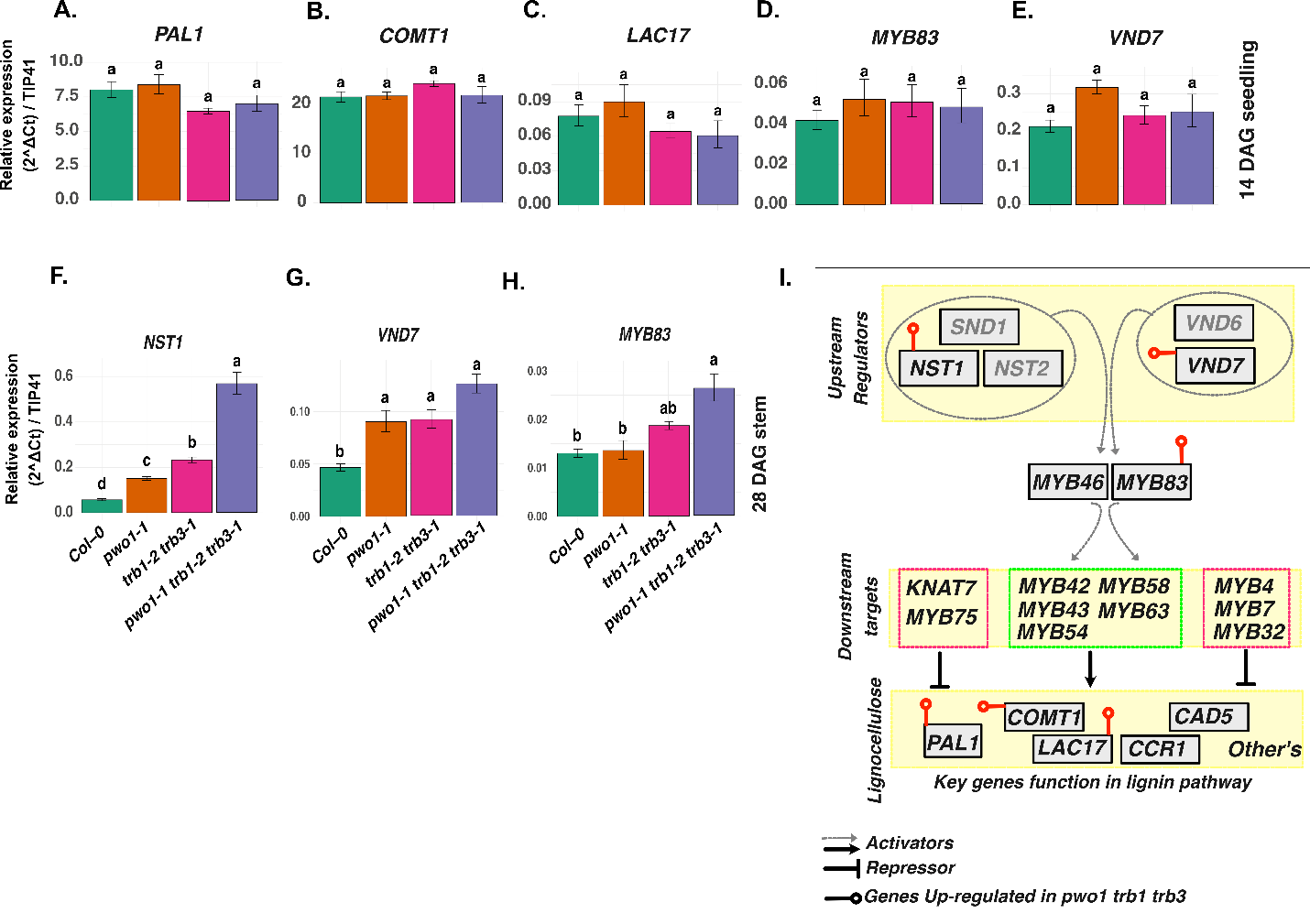

**Figure S14.** **qRT-PCR analysis of lignin biosynthesis and secondary cell wall genes.** Relative expression levels of lignin biosynthesis genes **A.** PAL1, **B.** COMT1, and **C.** LAC17, and secondary cell wall synthesis genes **D.** MYB83 and **E.** VND7 in 14-day-old seedlings. Expression of **F.** NST1, **G.** VND7, and **H.** MYB83 was also assessed in the main stems of 28-day-old plants after bolting and before flowering. **I.** Schematic representation of secondary cell wall biosynthesis showing key genes and regulators acting at upstream, intermediate, and downstream stages of the lignocellulose pathway. Red dumbbells mark genes significantly or consistently upregulated in the ***pwo1 trb1 trb3*** triple mutant compared with ***Col-0***, ***pwo1***, and ***trb1 trb3*** control plants. Data are mean ± SE from ≥3 biological replicates; different letters indicate significant differences by Tukey’s HSD test (*p* < 0.05).
